# Population-scale organization of cerebellar granule neuron signaling during a visuomotor behavior

**DOI:** 10.1101/200824

**Authors:** Sherika J.G. Sylvester, Melanie M. Lee, Alexandro Ramirez, Sukbin Lim, Mark S. Goldman, Emre R.F. Aksay

**Author notes:** Corresponding Authors: Mark S. Goldman, University of California at Davis, Davis, CA 95618; Emre R.F. Aksay, 1300 York Avenue, Weill Cornell Medicine, New York, New York 10021.

## Abstract

Granule cells at the input layer of the cerebellum comprise over half the neurons in the human brain and are thought to be critical for learning. However, little is known about granule neuron signaling at the population scale during behavior. We used calcium imaging in awake zebrafish during optokinetic behavior to record transgenically identified granule neurons throughout a cerebellar population. A significant fraction of the population was responsive at any given time. In contrast to core precerebellar populations, granule neuron responses were relatively heterogeneous, with variation in the degree of rectification and the balance of excitation versus inhibition. Functional correlations were strongest for nearby cells, with weak spatial gradients in the degree of rectification and excitation. These data open a new window upon cerebellar function and suggest granule layer signals represent elementary building blocks underrepresented in core sensorimotor pathways, thereby enabling the construction of novel patterns of activity for learning.

**SIGNIFICANCE STATEMENT:** Cerebellar processing is important for a variety of fine motor tasks and sensorimotor adaptations, and a growing body of evidence indicates a prominent role in cognitive control. However, it has been challenging to understand cerebellar function during behavior because of difficulties in recording from cerebellar granule neurons, the most populous neuron type in the brain. We use population-scale optical imaging in the larval zebrafish to compare precerebellar activity to granule cell signaling. Our results suggest a behaviorally relevant expansion of precerebellar signaling representations at the granule layer of the cerebellum.

## INTRODUCTION

Cerebellar processing is instrumental for a wide range of adaptive motor and cognitive behaviors, and cerebellar dysfunction is thought to contribute to diseases ranging from ataxia to autism. Investigations of cerebellar anatomy have revealed a highly conserved microcircuitry across species, with a pattern of feedback interactions with other brain regions suggestive of modular organization (Strick et al., 2009). This structural regularity may underlie general principles of cerebellar processing.

Common to all cerebellar circuits is the presence of a large granule cell population at the input layer. Granule neurons are the principal recipients of pre-cerebellar excitatory input via mossy fibers, are recipients of inputs from other cerebellar neurons including Golgi and unipolar brush cells, and are the primary source of excitatory signals to the cerebellar output layer via ascending and parallel fiber inputs to Purkinje neurons. The synapses between granule cells and Purkinje neurons can exhibit both long-term depression and long-term potentiation (Jorntell and Hansel, 2006), thus providing a possible source of plasticity underlying cerebellar involvement in learning. Prominent hypotheses have argued that such plasticity would be most effective if the granule cell population provided an expanded representation of sensorimotor information relevant for a particular behavior (Marr, 1969; Albus, 1971; Fujita, 1982). In this way, changing the strength of distinct parallel fiber synapses would drive specific behavioral adaptations. However, because of the small size, close packing, and large number of granule cells, it has been difficult to understand the large-scale organization of activity in the granule population during behavior.

Here we address this gap by measuring response properties throughout a granule cell population in awake larval zebrafish while animals perform a visuomotor behavior. We focus on the optokinetic response, a simple and well-studied behavior common to all vertebrates in which the eyes track large-field movements in the visual world. The core elements of this circuit have been elucidated over several decades of work (Masseck and Hoffmann, 2009b) (Figure 1a). During left/right optokinetic stimulation with velocity steps, the activity of direction-sensitive retinal ganglion cells are pooled in the pretectum to provide a representation of large-field stimulus velocity. Many pretectal neurons sensitive to movement in the horizontal plane fire in a reciprocal manner, increasing firing for stimulation in one direction, and decreasing firing for the other (Mustari and Fuchs, 1990; Kubo et al., 2014). These pretectal signals are relayed to second-order vestibular nuclei which, with participation of the velocity-storage neural integrator, provide an eye-velocity command signal directly onto motor neurons (Stahl and Simpson, 1995; Beck et al., 2006). This eye-velocity signal is also sent along a secondary path to motoneurons through the velocity-to-position neural integrator to generate eye-position commands (McFarland and Fuchs, 1992; Pastor et al., 1994). Together, these signals provide the appropriate phasic/tonic balance in motoneuron drive needed to generate an eye position profile tracking stimulus position. Adaptations in this drive during learning are mediated primarily by Purkinje output neurons of the cerebellum, which receive excitatory climbing fiber inputs from the inferior olive (not shown) and parallel fiber inputs from the cerebellar granule cells. Granule cells receive via mossy fibers signals originating from each of the non-motor populations forming the core sensorimotor pathway for optokinetic behavior (Masseck and Hoffmann, 2009b), including the pretectum, vestibular nuclei, and neural integrator populations. Whereas the activity of Purkinje cells and core pathway populations has been relatively well studied, little is known about the responses of the large granule cell population.

**Figure 1:**
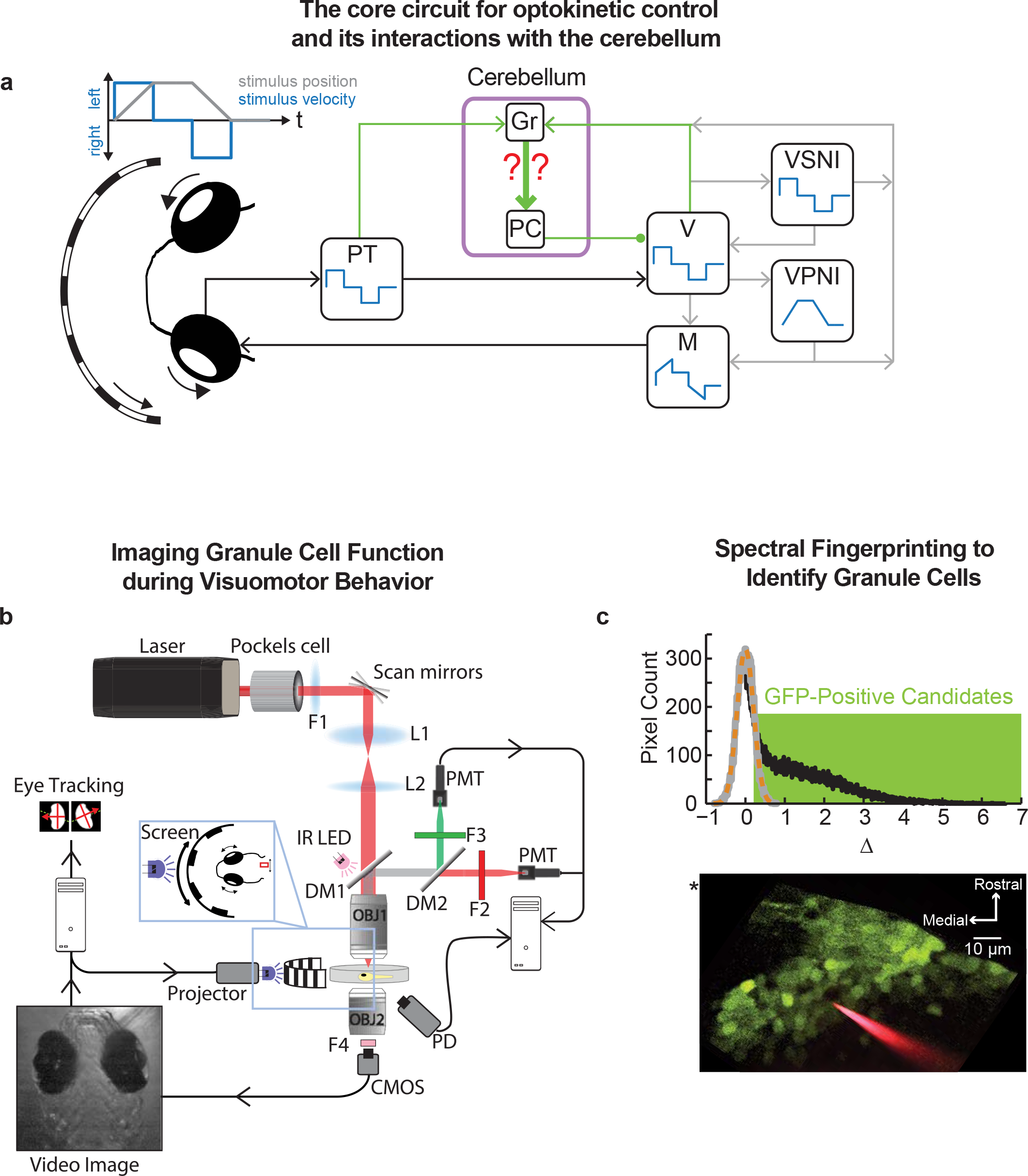
Imaging the signaling properties of transgenically identified granule cells during oculomotor control. (**a**) Current understanding of the signal flow underlying the control of optokinetic behavior. The core sensorimotor elements of the circuit are the pretectum (PT), second-order vestibular nuclei (V), the velocity-storage neural integrator (VSNI), and the velocity-to-position neural integrator (VPNI). Adaptations in oculomotor behavior are mediated by the Purkinje (PJ) and granule (Gr) cells of the cerebellum. In the mammal, pretectal signals and stimulus information processed in a cortical path are combined at the dorsolateral pontine nucleus before being sent to the cerebellum(Mustari et al., 1994). Filled circle-inhibitory connections; arrows – excitatory connections; for simplicity, callosal projections not explicitly indicated. (**b**) Experimental setup. Two-photon excitation with a Ti-Sapphire laser tuned to the infrared frequency range enables simultaneous fluorescence microscopy of granule cell function, behavioral stimulation, and eye-movement tracking. *Abbreviations* OBJ: objective, L: lens, F: filter, PMT: photomultiplier tube, PD: photodetector, DM: dichroic mirror, IR LED: infrared light emitting diode. (**c**) Spectral finger-printing procedure allowing identification of GFP-positive granule neurons. (Top) A histogram of the pixel-by-pixel differences in intensity (uint8) between two excitation wavelengths (Δ, black) was compared to the histogram of differences expected from noise (grey) (fit of noise histogram as dashed orange line). Pixels with difference values one noise standard deviation or more from zero were considered GFP-positive candidates (green box). (Bottom) Candidate GFP-positive pixels (green), at one image plane, identified in the inner granule layer (IGL) of the right lobe of the transgenic zebrafish cerebellum. Also visible is an electrode for glutamate iontophoresis (red). The location of the center of the midbrain-hindbrain border (*) is visible at the upper left.

We used two-photon calcium imaging with single-cell resolution to assess granule cell representations at the whole-population scale during optokinetic behavior. In the following, we describe activity throughout a granule cell population, the spatial organization of this activity, and how activity patterns differ from those in representative core pathway populations providing mossy fiber inputs. We also explore through a model implications of these data for involvement of the granule cell population in cerebellar learning.

## MATERIAL AND METHODS

### Zebrafish procedures for imaging granule neurons

#### Zebrafish preparation

All procedures were approved by the Institutional Animal Care and use Committee of Weill Cornell Medical College. Experiments were performed using Casper pigmentless mutants expressing the transgene *gata1:GFP* (strain 781) to allow unambiguous identification of cerebellar granule cells. In the cerebellum of *gata1:GFP* animals, cells with GFP expression are selectively colocalized with those neurons expressing the α6 subunit of the GABA_A_ receptor (Volkmann et al., 2008), an evolutionarily conserved subunit exclusively expressed at the cerebellar granule cell in all vertebrates (Kato, 1990; Bahn et al., 1996; Dino and Mugnaini, 2008). Larval zebrafish 5-8 days post-fertilization were prepared by first anesthetizing the fish in 100-200 mg/L ethyl 3-aminobenzoate methanesulfonate (E10521, Sigma) dissolved in 10mM HEPES (H3375, Sigma), and then embedding the fish in 1.7% agarose (A0701, Sigma) within the lid of a plastic petri dish (35-1008, BD Falcon) containing a layer of solidified 2% agarose (V3121, Invitrogen).

#### Indicator loading

Granule neuron activity in the inner granule layer (IGL) was monitored using the calcium-sensitive indicator Oregon Green BAPTA 1-AM (OGB) (O6807, Invitrogen). OGB was prepared by diluting to 5 mM with 10% (wt/vol) pluronic (P6867, Invitrogen) in dimethyl sulfoxide (276855, Sigma). OGB was delivered to the cerebellar corpus of anesthetized fish using five to ten 100-300 msec duration pulses (Picospritzer III, Parker Hannifin) through a borosilicate microcapillary (1B100F-4, WPI) with a tip diameter of 2–4 μm. Granule cells in the separate lateral (*eminentia granularis*) and caudal (*lobus caudalis*) subgroups were not reliably labeled with these injections and will be investigated in future work. Injections and subsequent experiments were performed in 10% (vol/vol) glucose-free Evans medium (in mM): 134 NaCl, 2.9 KCl, 2.1 CaCl_2_, 1.2 MgCl_2_, and 10 HEPES, pH 7.8. After injection, zebrafish were allowed at least four hours for dye-loading and recovery from anesthesia.

#### Fluorescence imaging

Fluorescence was monitored with a custom-designed two-photon microscope (Fig. 1b). The embedded zebrafish were placed under a water immersion 40x/0.8 numerical aperture objective (OBJ1; LUMPlanFl40XW/IR2, Olympus) that focused the excitation light from a MaiTai HP laser (Newport). Multiple horizontal planes through the cerebellum were imaged at a 2 Hz scan rate (512 × 512 pixels, ~3 planes per fish). Emitted light was collected through the 40x objective, and reflected by two dichroic mirrors (DM1, 720dcxruv; DM2, 565dcxr; Chroma Technology) into separate red (F2; ET605\70m-2p, Chroma) and green (F3; ET525\50m-2p; Chroma) channels. Fluorescence signals were detected and amplified by photomultiplier tubes (PMTs, R3896; Hamamatsu), while scattered light from the focal plane was detected by a substage photodetector (PD, PDA36A, Thorlabs). Signals from the PMTs and the PD were acquired by ScanImage(Pologruto et al., 2003) to generate fluorescence and contrast images of specimen in the focal plane.

#### Behavior

We utilized the robust optokinetic responses of the larval zebrafish(Beck et al., 2004) to probe the functional properties of cerebellar granule neurons in the awake preparation. To allow tracking of eye movements and an unobstructed view of the optokinetic stimulus, wedge-shaped agar pieces were removed from in front of the eyes. The zebrafish was illuminated from above by an infrared LED (IR LED; LED851L, Thorlabs) placed near the back face of the first objective (OBJ1). A second objective (OBJ2, 4X/0.1 NA, WPI) was positioned below the zebrafish to create an image detected by a CMOS camera (F036B, Allied Vision Technologies). The camera was protected from laser light by two filters (FB850-40, Thorlabs; F4, XF3308, Omega). Eye movements were calculated in custom Matlab-based software as previously described(Miri et al., 2011a). The optokinetic reflex was evoked by the horizontal movement of a computer-generated image in a sequence of velocity steps with an amplitude of 8°/sec and a duration of 8 or 20 seconds. The image, composed of vertical bars 1 cm wide and 1 cm apart, was projected (MPro110, 3M) onto a 1 cm tall and 6 cm wide screen (NT4008, Novatron) placed approximately 2 cm in front of the fish (Fig. 1b, inset). Two filters (FGB25 & FES0450, Thorlabs) were placed in front of the projector to limit interference with fluorescence detection.

### Imaging granule cell activity during oculomotor function

#### Motion Correction

Zebrafish at times made head or body movements that generated a few micrometers of image displacement. These global changes in position were corrected following a semi-automated procedure by aligning all image frames of a recording to a reference image (Miri et al., 2011a). In the first step of correction, the reference image was an average of hand selected frames where, qualitatively, little movement was evident. Image frames where the shift was in excess of a threshold 0.15 μm were then excluded from further analysis. For the second iteration, the reference image was an average of all remaining image frames, and frames were again excluded if the new shift was greater than the threshold. Shift-corrected movies were then inspected and blocks were excluded from further analysis if they contained a large number of eliminated frames. On average, a total of 22.10% of the frames were excluded from analysis.

#### Granule cell identification

Identification of granule cells in the inner granule layer (IGL) after indicator loading was achieved by a simple “spectral fingerprinting” procedure that exploited the bimodal peak excitation of OGB around 790 nm and 930 nm and the unimodal peak excitation of GFP around 930nm(Brondi et al., 2012) (Fig. 1c). Fluorescence recordings of optokinetic responses at each image plane were collected for identical periods of 3-5 minutes at the two excitation wavelengths, at the same power. At each wavelength *λ*, the time-averaged fluorescence intensity at a particular pixel *i*, *F̅*_*λ,i*_, is given by

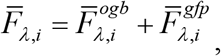

where 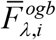 and 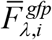 are the contributions from the two fluorophores. For each fluorophore, the average fluorescence values contributed at the two wavelengths are related by a scale factor *a*. Thus

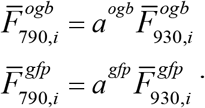

To identify those pixels corresponding to GFP-positive granule cells, we calculated a difference, Δ_*i*_, that eliminated the contribution from OGB fluorescence:

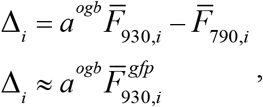

where the second line follows by substitution and the fact that, because of the differences in excitation spectra, *a*^*ogb*^ ≫ *a*^*gfp*^. To determine the scale factor *a*^*ogb*^, we took advantage of the fact that the adjacent crista cerebellaris, which we found in control animals were absent of GFP-positive soma, were well-labeled by OGB. Thus, the scale factor was calculated by taking the ratio (790/930) of time-and pixel-averaged fluorescence values determined from a ~50 micrometer diameter region in the crista. After this procedure, difference data for GFP-negative cells can be expected to have mean of zero, while data for GFP-positive cells should be positive (Fig. 1c, top, black).

To identify a threshold for assigning pixels to either the GFP-negative or GFP-positive group, we fit a Gaussian curve (Fig. 1c, top, dashed) to a mirror-symmetric sample generated from the portion of the histogram below 0 (top, gray). The standard deviation of this Gaussian established the threshold, with Δ_*i*_ values below this threshold set to zero, and those above threshold taken as candidates for inclusion (top, green). We next identified contiguous regions containing more than 100 pixels where Δ_*i*_ values were greater than threshold using Matlab’s “bwboundaries” function (bottom). Regions of Interest (ROI) specifying individual GFP-positive granule cells of 2-4 μm diameter were then drawn manually through a user-interface (with Matlab’s “roipoly” as the main component) using both this Δ- map and the average 930 nm image. As a final confirmation of GFP expression, the number of pixels within each user-drawn ROI that overlapped the GFP-positive pixel map were counted, and ROIs with areas that were at least 75% GFP-positive were used for the remainder of this study.

#### Fluorescence Time Series

The change in OGB intensity for each granule neuron was computed using the ROIs drawn during the spectral fingerprinting procedure. For each frame collected at 930 nm, fluorescence intensity was averaged across all pixels bounded by an ROI. The fluorescence time series of a given cell was determined as the percent change in fluorescence intensity with respect to the average fluorescence intensity (which included contributions from both OGB and GFP). Each fluorescence time series was interpolated at 20 Hz and then averaged across each cycle to generate an optokinetic cycle-triggered average used during analysis.

### Interpreting granule fluorescence signals

#### CIRF time constant estimation

To quantitatively interpret fluorescence signals, it was necessary to first determine the calcium impulse-response function (CIRF) of granule neurons. The CIRF is a measure of the intracellular calcium dynamics associated with a brief synaptic excitation or the firing of a brief burst of action potentials; once known, it can be used to either deconvolve fluorescence signals to determine changes in electrophysiological responses (Yaksi and Friedrich, 2006), or to convolve a model of changes in electrophysiological response and fit to fluorescence data (Miri et al., 2011a). Here we measured the CIRF of granule neurons by monitoring fluorescence changes triggered by localized iontophoretic delivery of glutamate.

Iontophoretic stimulation of granule neurons was accomplished by delivering brief pulses of glutamate through a targeting electrode. First, the skin was perforated under anesthesia near the midbrain-hindbrain border by a microcapillary coated with DiI crystals. Upon recovery from anesthesia, an electrode with a sub-micron diameter tip filled with a 100 mM glutamate (2469-57, Mallinckrodt Chemicals) and 0.5 mM Alexa 594 salt (A-10438, Invitrogen) in 100% Evans solution was guided under fluorescence imaging through the perforation so that the electrode tip was placed near GFP-positive granule cells (Fig. 1c, bottom, red; 15-20MΩ). A ground electrode was also placed in the surrounding bath. For iontophoresis, trains of −5 to −8 V, 2 msec electrical pulses were administered at 200 Hz for 0.5 sec once every 20 seconds (Fig. 2a, top). Through trial and error, we found that these settings were the minimum needed to elicit consistent responses; larger amplitude stimulations were also effective, but generally resulted in longer-lasting responses indicative of either tonically elevated extracellular glutamate or reverberation. The CIRF time constants following minimal stimulation were measured by exponential fits (top left, orange) to stimulus-triggered averages (top left, black).

**Figure 2:**
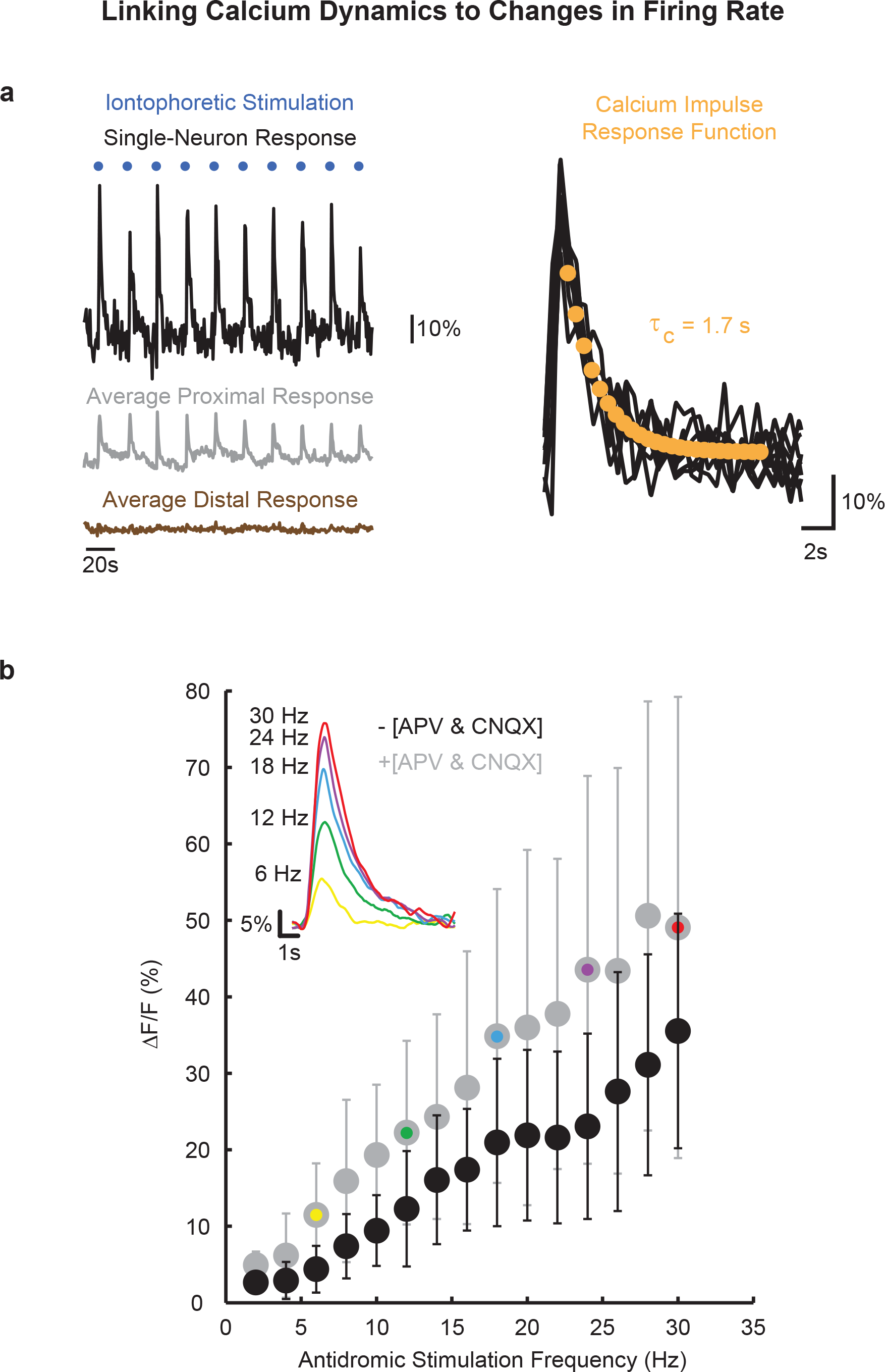
Links between changes in calcium concentration and firing rate in granule neurons. (**a**) To assess the calcium-buffering dynamics of identified granule neurons, glutamate pulses were administered via iontophoresis into the IGL every 20 sec (left, blue dots) to elicit a brief change in local granule neuron activity. The fluorescence response of one granule cell is shown in black, the average response of those neurons within 5 μm of the electrode tip is shown in gray, and the average response of cells between 5 and 20 μm from the tip is shown in brown. Pulse-triggered averages for 10 activated granule neurons (right, black) were fit with an exponentially decaying function (the Calcium Impulse Response Function, or CIRF) to estimate the time constant of buffering ( *τ*_*c*_, orange).(**b**) Granule neurons were antidromically activated by depolarizing pulses administered to parallel fibers in the contralateral molecular layer in the absence (black) and presence (gray) of APV and CNQX. The responses of active granule neurons were averaged and the peak average response is plotted (gray: 2 fish, 12 cells; black: 2 fish, 16 cells). The average response of one cell for 6, 12, 18, 24, and 30 Hz are displayed in the inset (horizontal bar: 1 second, vertical bar: 5%). Each colored dot corresponds to an average response in the inset.

#### Assessing the relationship between firing rate and calcium signal

To assess the linearity of the relationship between changes in firing rate and calcium dynamics, we examined the somatic calcium dynamics of granule neurons in response to antidromic stimulation at different frequencies. To trigger action potentials, a stimulus current was administered by an electrode containing 1x Evans solution and 0.3mM Texas Red (D-3329, Life Technologies) at segments of the caudal lobe parallel fiber commissure that were located contralateral to the site of OGB recording (Volkmann et al., 2008). To prevent an excess amount of the electrode solution from flooding the ventricular space, the tip of the electrode was front-loaded with molten 1.7% (m/v) agarose (A0701, Sigma) that was allowed to solidify before backloading with the Texas Red solution. The parallel fiber commissure was stimulated every 10 seconds over a span of 1 minute by a 1 second long pulse train (1 msec per pulse) with a frequency ranging from 2 Hz to 30 Hz and a voltage of −20 V (Fig. 2b). GFP-positive neurons activated by antidromic pulses were first identified by heightened changes in fluorescence following 30 Hz pulses. Once a pocket of activated cells was identified, the frequency of the stimulus was dialed down by increments of 2 Hz for each subsequent stimulation sequence. To check for continued electrical discharge of affected parallel fibers and cell responsiveness, an additional set of 30 Hz electrical pulses were administered following the conclusion of the 30 to 2 Hz stimulation sequence. Neurons were selected for further analysis if their average peak fluorescence values exceeded 5% during the 30 Hz experimental and post-experimental stimulations, and their average fluorescence response during 10 and 30 Hz stimulations were highly correlated (≥ 0.7) to a response template created by averaging post-30 Hz stimulation responses with amplitudes greater than 50%.

To determine the origin of somatic calcium changes, we compared somatic calcium responses of synaptically active and inactive granule neurons (separate populations). Inactivation of NMDA and AMPA receptors was achieved by injecting a solution containing 1mM APV (A-8054, Sigma), 1mM CNQX (C-239, Sigma), and 0.3mM Texas Red directly into the hindbrain within proximity of the *crista cerebellaris*. The synaptic blocker solution was allowed to diffuse throughout the entire brain for 10 minutes prior to antidromic stimulation and calcium imaging as above.

#### Quantifying granule cell activity

We next fit granule neuron data to a model of expected fluorescence variation. As shown in the Results (Fig. 3), the relative responses to ipsiversive vs. contraversive stimulation varied greatly across granule neurons. To capture this variability, we defined for neuron *i* a velocity-sensitivity profile *Vi(t)* such that

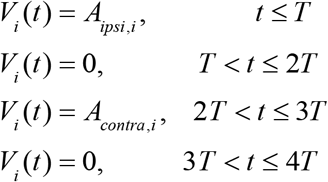

where *A*_*ipsi*_ and *A*_*contra*_ give the sensitivity to ipsilaterally- and contralaterally-directed stimulus movement, and *T* refers to the duration of one stimulus epoch (Fig. 3a bottom, blue). Additionally, we noted that the responses of many neurons were more consistent with a low-pass filtered velocity signal (*e.g.* cell 2 in Fig. 3a). Therefore, the electrophysiological response *r* for granule neuron *i* was specified by the convolution

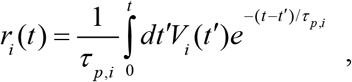

where *τ*_*p*_ is a free parameter characterizing a granule cell’s persistence time constant (Fig. 3a bottom, purple). We also note that because saccadic or fast-phase eye movements were infrequent in these data and the granule cell responses to these events appeared modest at best, these inputs were not included in the model.

**Figure 3:**
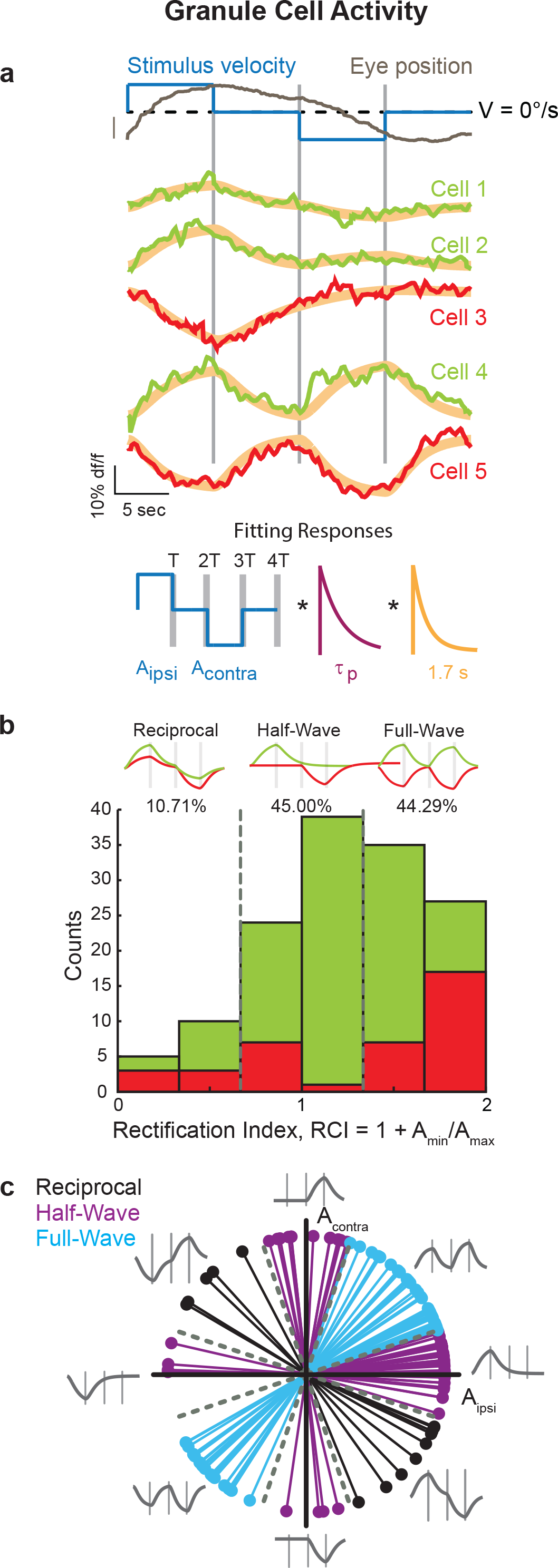
Optokinetic responses of cerebellar granule neurons. (**a**) Top: In the inner granule layer (IGL), steps in stimulus velocity (blue) drove optokinetic response (dark gray) that revealed heterogenous activity patterns. Representative reciprocal (cell 1), half-wave (cells 2 and 3), and full-wave (cells 4 and 5) rectified cycle-triggered fluorescence responses; green indicates positive, and red indicates negative rectification. Not all data from the same image plane. Gray bars denote the time of change of stimulus velocity. Model fits are in orange. Scale bar: 5°. Bottom: Fluorescence responses were modeled as encoding a low-pass filtered optokinetic velocity signal that then gets filtered through a calcium impulse response function that models the kinetics of calcium buffering. Under this model, the applied optokinetic velocity signals (blue) were convolved first with a low-pass (exponential decay) filter of time constant *τ*_*p*_ (maroon) and then with a second low-pass (exponential decay) filter of time constant *τ*_*c*_ representing the effects of intrinsic calcium buffering (orange). The time constant *τ*_*c*_ was empirically determined via glutamate stimulations (see CIRF time constant estimation section) and the time constant *τ*_*p*_ was fit to the fluorescence and applied optokinetic velocity traces. (**b**) Histogram of the number of neurons within each response type. Grey dashed lines at rectification index (RCI) values of 2/3 and 4/3 denote response type boundaries. Green indicates positive and red indicates negative rectification for half-and full-wave responses; for reciprocal cells, the two colors indicate whether the positive (green) or negative (red) epoch achieved the greatest magnitude. (Top) Model fits to typical neurons in each subgroup with ipsiversive movement sensitivity (i.e. excitatory response to ipsiversive movement and/or inhibitory response to contraversive movement); those with contraversive movement sensitivity are also included in the histogram. (**c**) For granule neurons, polar plot indicating the relative contribution of the fit coefficients A_ipsi_ and A_contra_. Grey dashed lines denote the RCI response group cutoffs in polar coordinates.

Next, to directly fit this model to the fluorescence data, the electrophysiological response model was convolved by the CIRF to generate a model of the fluorescence time series *F(t)* for cell *i*,

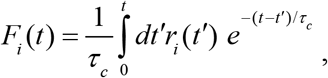

where *τ*_*c*_ is the population-average CIRF time-constant (Fig. 2a, bottom; Fig. 3a bottom, orange). Nonlinear regression of this model to the cycle-triggered average fluorescence data was performed using the “nlinfit” function in Matlab to determine the unknown parameters *A*_*ipsi*_, *A*_*contra*_, and *τ*_*p*_ for each neuron. The *τ*_*p*_ values were capped at 25 seconds when reporting population statistics.

Quantitative response measures were determined for a subset of neurons with high signal-to-noise ratios. A GFP-positive cell was selected for further characterization if its fit r-squared value was above 0.5, its signal to noise ratio, determined as the amplitude of the fit divided by the standard deviation of the residual, was greater than 3, and the amplitude of its maximum response was greater than or equal to 5% ΔF/F.

We evaluated several response measures for these well-fit, high signal-to-noise data. Because such fluorescence measurements can only be used to assess relative changes in activity, measures were based on comparing response coefficients across stimulation periods. First, for each cell a rectification index (*RCI*) was specified as 1+*A*_*min*_/*A*_*max*_ where *A*_*min*_ and *A*_*max*_ are the velocity-sensitivity coefficients whose magnitudes are the smallest and the largest, respectively, regardless of their association with ipsiversive or contraversive stimulus movement. Second, data were characterized through a polar plot in which each granule neuron response was described by its relative response to ipsiversive and contraversive stimulus velocities as θ = tan^−1^(*A*_*contra*_/*A*_*ipsi*_). Data with (*A*_*ipsi*_ *A*_*contra*_) values that were (+ +), (− +), (− −), or (+ −) were assigned to quadrants 1-4, respectively. Third, a signed rectification index (RSI) was determined by *RCI*\**sign(A_max_)* to assess the possible contribution of excitatory vs inhibitory inputs. Fourth, a direction-sensitivity index (*DSI*) defined as (*A*_*ipsi*_ − *A*_*contra*_) / |*A*_*max*_| was used to assess ipsiversive vs contraversive sensitivity.

The spatial organization in the *RCI*, *RSI*, and *DSI* activity indexes was determined in two ways. First, we pooled information on cell locations across all fish after registration to the midline and midbrain-hindbrain border, then fit a plane to the data in the space composed of cell activity index and location coordinates. Second, to minimize the effects of possible nonlinearities and registration errors, we calculated for every pair of granule cells within a given fish the pairwise difference in activity index (*RCI*, *RSI*, *DSI*) and the pairwise distance, then pooled data across fish. In this case, spatial trends were assessed using the Spearman Rank Correlation (src), and statistical significance was determined using the two-tailed Student’s t-test. Data were grouped into 20 bins with equal sample numbers, and then the mean +/− s.e.m. index difference was plotted against the mean separation distance for each bin. Trends from pairwise measures were assessed along the rostro-caudal and medio-lateral axes, as well as the major and minor axes of the IGL population determined from fitting an ellipse to the pooled cell location data. Sensitivity to outliers was checked using a bootstrap procedure (sampling with replacement, 1000 iterations) and robust regression (Matlab *robustfit*, bisquare, 4.685 tuning constant).

### Identification and imaging of core sensorimotor pathway neurons

#### Preparation

To better understand how granule neuron activity related to the signals along the core optokinetic sensorimotor pathway, we also recorded the activity of hindbrain sensorimotor pathway neurons during velocity steps. These recordings included vestibular neurons, the velocity-storage neural integrator (VSNI), a cell group that provides velocity-related signals during optokinetic tracking (Pastor et al., 1994; Beck et al., 2006), and the velocity-to-position neural integrator (VPNI), a cell group responsible for generating position signals(Lopez-Barneo et al., 1982; McFarland and Fuchs, 1992; Aksay et al., 2007; Miri et al., 2011b) (Fig. 1a). Vestibular neurons were recorded in rhombomeres 3-6 (within 30 microns dorsal or ventral of the medial longitudinal fasciculus), in accordance with prior anatomical studies (Straka et al., 2001; Matsui et al., 2014); the VSNI and VPNI populations were recorded in rhombomeres 7-8, in accordance with prior anatomical and functional findings (Miri et al., 2011b; Daie et al., 2015). Experiments recording the VSNI and VPNI populations were conducted using 5-8 days post-fertilization zebrafish carrying the transgene *vglut2:dsRed* (Kinkhabwala et al., 2011), a line in which glutamatergic neurons are well-labeled. The procedures for zebrafish and OGB preparation were as described above. OGB was injected in rhombomeres 7 and 8, approximately 150μm below the skin, in the lateral portion of the caudal hindbrain. After several hours of recovery, the fluorescence activity of neurons at various depths in rhombomeres 7 and 8 was recorded during optokinetic stimulation as described above. Vestibular neurons were recorded in 6-8 dpf animals from the *Tg(elavl3:H2B-GCaMP6f)* line (Dunn et al., 2016) as these populations were more difficult to load using OGB.

#### Identification of neurons

Cells with activity patterns related to oculomotor parameters were identified using a modification of a semi-automated strategy based on pixel-level correlations(Miri et al., 2011a). Here, to allow identification of cells with varying degrees of signal rectification, correlations to half-wave and full-wave rectified stimulus and eye movement parameters were also determined. Regions of Interest were drawn on a maximal-correlation map. Putative inferior olivary neurons, identified by their close-packing and extreme ventro-caudal location, were excluded from the current analysis and will be considered in future work. Neuronal activity was then fit with the same procedures described above for granule neurons; the CIRF time constant, measured previously for VPNI neurons, was set to 1.9 seconds (Miri et al., 2011b).

### Experimental design and statistical analysis

All statistical design and analyses are reported in the Results. In brief, differences in the stimulation responses presented in Figure 2B were assessed using the two-sample Kolmogorov-Smirnov (KS) test. Differences in distribution of rectification values between granule and precerebellar populations were also assessed with the KS test; differences in the mean relaxation time constant were assessed with the two-sample T-test. Spatial patterns in functional indices (Fig. 7) were assessed using the Spearman Rank Correlation (src). All tests were performed using Matlab.

## RESULTS

We report on activity throughout a granule cell population during optokinetic behavior. In the following, we first present the key aspects of our recording approach and elucidate the links between changes in calcium concentration and the changes in firing rate of granule cells. Second, we describe the functional properties during optokinetic stimulation of zebrafish granule neurons. Third, to better place these granule neuron patterns into context, we also determine the activity patterns under identical stimulus conditions in three of the core sensorimotor pathway populations involved in optokinetic behavior. Fourth, we investigate the spatial organization of granule cell activity. Finally, we use a phenomenological model to consider how the observed granule cell coding properties might contribute to the cerebellar control of learning.

### Calcium imaging throughout a granule cell population

The functional properties of cerebellar granule cells in the larval zebrafish were recorded during optokinetic tracking behavior (n = 6 fish). Recordings were obtained throughout the inner granule layer (IGL) of the cerebellar corpus (Nieuwenhuys, 1967; Bae et al., 2009), the most populous of the three granule layer subdivisions in teleosts, while zebrafish performed horizontal optokinetic tracking. To record granule cell activity, we extended our previously described procedures for calcium imaging in the awake zebrafish preparation (Miri et al., 2011a; Daie et al., 2015) by coupling two-photon fluorescence microscopy with optokinetic stimulation and video-tracking of eye movements (Fig. 1b). Optokinetic response activity was monitored at the single-cell level in one half of the cerebellum at different horizontal planes throughout the IGL after bolus-loading of the high-affinity calcium sensor Oregon Green BAPTA-1 AM (OGB). To distinguish granule cells from other cerebellar neurons also loaded with indicator dye, we used the *gata1:GFP* transgenic line, where neuronal expression in the cerebellum is limited to granule cells (Volkmann et al., 2008). GFP-positive neurons were identified through excitation fingerprinting by comparing fluorescence intensities at two wavelengths, one at which primarily OGB was excited, and another at which both OGB and GFP were excited (Fig. 1c; Materials and Methods). Through this procedure, 543 indicator-loaded granule cells were identified in the IGL; these were located within 100 μm of the dorsal surface, within 120 μm of the midline, and within 80 μm caudal of the center of the midbrain-hindbrain border (cMHB, * in lower panel of Fig. 1c). The cells were almost evenly distributed across fish (n = 94, 88, 69, 116, 106, and 70 for fish 1-6, respectively; average of 90.5±18.9 cells from each animal). For comparison, we found that in individual fish there were approximately 700 GFP-positive IGL neurons per lobe (in comparison to 50-100 Purkinje cells (Kaslin et al., 2013; Hamling et al., 2015)). Given the high degree of coverage in this genetic line (Volkmann et al., 2008), this sampling likely provided a thorough evaluation of the IGL population.

To guide our analysis of fluorescence responses during optokinetic behavior, we first characterized in three ways the association between calcium dynamics and changes in the firing rate of granule cells. First, we measured changes in granule neuron fluorescence accompanying brief stimulation with localized glutamate iontophoresis mimicking a burst of mossy fiber input. Responses rose rapidly during stimulation, and following stimulation returned to baseline in an exponential fashion with a time constant *τ*_*c*_ that had a mean and standard deviation of 1.66 +/−0.25 seconds, and a range from 1.17 to 1.99 seconds (n=10 cells; Fig. 2a). This relaxation timescale likely reflects the slow clearance of calcium following cessation of action potential discharge, as suggested by *in vitro* work on granule cells (Gandolfi et al., 2014) and *in vivo* studies of other cell types involved in optokinetic processing in the zebrafish (Miri et al., 2011b; Daie et al., 2015). In the following we refer to this exponential function with a 1.7 second time constant as the calcium impulse response function (CIRF, *τ*_*c*_) for granule cells.

Next, we investigated the linearity of changes in fluorescence with known changes in firing rate. Action potentials were triggered by electrically stimulating parallel fibers in the caudal molecular layer of the contralateral half of the cerebellum, targeting a location from which we could retrogradely label neurons in both the IGL and *eminentia granularis* (EGL, a more lateral granule cell subpopulation). The peak value of average fluorescence change in granule somata increased in a linear manner over the 2 Hz to 30 Hz stimulus frequency range to a maximum of around 35% (Fig. 2b; black, r^2^ = 0.98, n=12 cells). Additionally, the average relaxation of fluorescence following the peak was well fit by an exponential with the 1.7 s time constant determined above (r^2^ = 0.68 ± 0.27 for frequencies above 5 Hz; data not shown). Together, these results suggest that we can translate between firing rate and changes in calcium concentration over a wide dynamic range using a simple linear filter model. Such a model for interpreting fluorescence changes during optokinetic behavior is developed in the next section.

Finally, we investigated what sources might be dominating calcium influx at the soma. Both high-threshold voltage-sensitive calcium channels (VSCC) and voltage-sensitive N-methyl-D-aspartate receptors (NMDAR) provide significant calcium influx into granule cells (with calcium changes potentially amplified by calcium-induced calcium release from internal stores). Direct influx of calcium into the soma is expected through somatic VSCC activated during action potential discharge (D'Angelo et al., 1998; Gandolfi et al., 2014). Indirect influx to the soma might arise through diffusion of calcium from the dendritic terminals. Calcium accumulates in the dendritic terminals of granule cells in two ways: one, through the opening of dendritic VSCC with action potentials (because the small granule cells are effectively isopotential), and two, through dendritic NMDAR activation by combined mossy-fiber input and action-potential discharge (D'Angelo et al., 1994; Gall et al., 2005; Schwartz et al., 2012) (because of the voltage sensitivity of NMDA receptors). To examine the possibility of an indirect calcium influx, we therefore repeated the antidromic stimulation experiments in the presence of synaptic blockers APV (NMDAR blocker) and CNQX (AMPA receptor blocker). We expected that if indirect influx played a significant role, that we would observe antidromic somatic responses that would be significantly smaller or negligible since the NMDA-mediated component would now have been eliminated. We found that responses at each stimulation frequency were comparable or slightly larger than observed in the absence of blockers, and generally did not differ significantly (Figure 2b; gray, n=16 cells; p < 0.05 for all stimulation frequencies except 2, 6, 8, and 10 Hz; Two-Sample Kolmogorov-Smirnov (KS) Test). The overall trend in the responses was again linear over a wide range (gray, r^2^ = 0.99; inset shows average responses at selected frequencies for one cell), and responses were again well-fit by the 1.7 second exponential (r^2^ = 0.92 ± 0.05 for frequencies above 5 Hz). Together, these data suggest that there is little indirect influx of calcium to the soma, and that changes in somatic calcium recorded *in vivo* stem from influx through somatic voltage-sensitive calcium channels, in agreement with *in vitro* studies suggesting a spatial localization of calcium dynamics in granule cells(Gall et al., 2005; Gandolfi et al., 2014).

### Functional properties of granule cells during optokinetic behavior

The signaling properties of granule neurons were recorded during horizontal optokinetic tracking of vertical bars moving back and forth with a profile composed of velocity steps (Fig. 3a–c). A cycle of the stimulus sequence consisted of movement in the ipsiversive direction at a constant speed, followed by a fixation period, and then stimulus movement in the contraversive direction at the same speed, followed by another fixation period (blue line). Optokinetic behavioral responses consisted of gradual changes in eye position during stimulus movement, with slow centripetal drift in eye position during the fixation periods (dark grey line; gain ~ 0.1). Intermittent spontaneous saccades and fast-phase responses (a rapid re-centering of the eyes during optokinetic stimulation) were also present (not shown), albeit less commonly so than has been reported previously (Beck et al., 2004).

We observed a range of granule cell activity patterns during optokinetic stimulation (Fig. 3a top). Generally, these could be well characterized in relation to a low-pass filtered version of the velocity profile of the stimulus, as activity typically relaxed slowly back to baseline values during epochs where the stimulus velocity was zero. A small minority of granule cells exhibited an increase in activity for one direction of motion and a reciprocal decrease in activity for the opposite direction (cell 1). Much more commonly, granule cells exhibited nonlinear relationships to stimulus movement, with varying degrees of signal rectification. In one such group (cells 2 and 3), activity changed during stimulus movement in one direction, but there was little or no change in activity for opposing movements. This group was subdivided into cells showing either an increase or a decrease in activity, which we referred to as either positive (cell 2) or negative (cell 3) half-wave rectification. In another group, fluorescence levels either increased or decreased during stimulus movement in a manner irrespective of direction; these neurons were described as exhibiting either positive (cell 4) or negative (cell 5) full-wave rectification. Saccade-related responses were minimal or absent (not shown).

To quantify these optokinetic responses, we performed a fit of the fluorescence time series to a model for granule neuron activity (Fig. 3a bottom; Materials and Methods). In the model, firing rate responses were specified by ipsiversive and contraversive velocity sensitivities*, Aipsi* and *Acontra*, and a low-pass filter time constant of value *τ*_*p*_. Model responses were then convolved by the empirically determined CIRF capturing the dynamics of calcium buffering. Note that these two time constants capture different physiological properties: *p* captures the possibility of dynamics in neuronal firing that persist at a longer time scale than the dynamics of sensory input, and thus likely reflects more motor or integrative aspects of the activity; *c* reflects the dynamics of calcium buffering associated with individual action potentials, and was determined by the measurements of Figure 2a.

Using the fitting procedure described, we found that 187 of the 543 granule neurons displayed changes in fluorescence that were well fit by the model (r^2^ ≥ 0.5; example fits shown in Figure 3a, orange). The average range of fluorescence responses for these cells was 9.62 ± 5.67%, within the dynamical range explored in the stimulation experiments of Figure 2. We also examined the residuals from the fits to determine if the model used was sufficient to capture the signal present in the data. We calculated the autocorrelation of the residual and determined the strength of the autocorrelation within a +/− 1 second lag, an indicator of remaining signal in the residual. We found that residual signal strength for cells that were well fit (r^2^ ≥ 0.5) was generally greater than those that were poorly fit (r^2^ <0.5) by the model (p = 4.19 × 10^−12^, Two-Sample T-Test). This suggests the model form used captured the signal present in the data. In the following, we proceeded to quantify response metrics for those cells with moderate to large response amplitudes (Materials and Methods, n = 140 cells).

We next used these fits to quantify activity patterns across the population. Given the qualitative disparity in the degree of rectification across cells (Figure 3a), we first defined a rectification index (*RCI* = *A*_*min*_/*A*_*max*_ + *1*), where **A*_*min*_* and **A*_*max*_* is assigned to the velocity sensitivity parameter with the smallest or largest magnitude, respectively. This index was formulated such that a perfectly reciprocal response would be given an *RCI* value of 0, a perfectly half-wave rectified response would have an *RCI* value of 1, and a full-wave rectified response with equal sensitivity to ipsilateral and contralateral stimulation would have an *RCI* value of 2 (see top of Fig. 3b for examples). For the granule cell population, a histogram of these *RCI* values revealed a distribution heavily skewed towards rectified responses (Fig. 3b). Similar trends were noted for neurons with positive (green, 72.86%) and negative (red, 27.44%) maximal response amplitudes. Very few granule neurons showed strongly reciprocal activity with *RCI* values of less than 1/3 (3.57%), and only 10.71% of cells had *RCI* values of less than 2/3. Neurons with *RCI* values between 2/3 and 4/3 reflecting a half-wave rectified waveform composed 45.00% of the population. Neurons with *RCI* values exceeding 4/3 reflecting a full-wave rectified waveform composed 44.29% of the cells. Neurons with strong full-wave rectification, with *RCI* values exceeding 5/3, were 19.29% of the total population, and were thus observed over 5 times as often as those cells with strong reciprocal activity (*RCI* values less than 1/3).

Neuronal responses were also quantified using a polar analysis to provide more granularity (Fig. 3c). An angular representation of rectification in each cell was determined by the inverse tangent of the ratio between the response-amplitude coefficients during back and forth image motion, *A*_*ipsi*_/*A*_*contra*_. Thus, neurons with reciprocal waveforms fell along the line with slope −1 (black), cells with half-wave rectified waveforms along the axes (purple), and neurons with full-wave rectified waveforms along the line with slope 1 (cyan). Weighting towards full-wave rectification in the granule cell population was reflected in the relative abundance of granule neuron data falling along the slope 1 line. Directional sensitivity was mixed, with 68.57% of neurons with reciprocal and half-wave rectified waveforms responding most strongly with ipsiversive movements.

The above analyses show that the functional properties of granule cells in the IGL during optokinetic tracking behavior are varied, covering the full spectrum of possible rectification values, although with relatively few reciprocal cells. Such high levels of rectification have not been reported in the core sensorimotor pathway neurons providing optokinetic-related mossy-fiber inputs, the pretectal (Mustari and Fuchs, 1990; Masseck and Hoffmann, 2009a; Kubo et al., 2014), vestibular (Stahl and Simpson, 1995; Portugues et al., 2014), and the VSNI (Pastor et al., 1994; Beck et al., 2006) and VPNI(McFarland and Fuchs, 1992; Pastor et al., 1994; Daie et al., 2015) populations, where reciprocal and positive half-wave rectified responses predominate (see also Discussion). To check that this rectification was not somehow an artifact of our methodology and approach, we next imaged responses during identical stimulus conditions in the two NI populations and the vestibular complex. The NI populations were a focus for two reasons: first, previous work suggests these form the largest group of responsive neurons during optokinetic stimulation (Portugues et al., 2014), and these populations provide a prominent source of mossy fiber input to the cerebellum (Straka et al., 2006; Ma et al., 2009). In a separate set of experiments, OGB was injected into the caudal hindbrain, and fluorescence changes of NI cells were monitored during back and forth optokinetic stimulation (Fig. 4a). Consistent with data from other species (McFarland and Fuchs, 1992; Pastor et al., 1994; Beck et al., 2006), and in contrast to granule cell signals, the vast majority were reciprocal (top two cells) or positive half-wave rectified (bottom two cells). These responses were also quantified with the fitting procedure described above (85 cells, 2 fish). Overall, the distribution of *RCI* values for NI cells was significantly different from that of the granule cell population (Fig. 4b; p = 3.71×10^−16^, KS Test). Only 15.29% of neurons had maximal responses that were negative, and only 4 of these were non-reciprocal cells. 34.11% of NI neurons displayed reciprocal responses, 61.18% had half-wave rectified responses, and only 4.71% were categorized as full-wave rectified; furthermore, no responses were found to be strongly full-wave rectified (*RCI* greater than 5/3). A polar plot of these responses showed that the NI population had predominantly ipsiversive sensitivity, and reaffirmed the stark contrast relative to the granule cell population in the degree of negativity and rectification (Fig. 4c). Similar results were found when comparing the granule population to the vestibular population (p = 4.34×10^−7^, KS Test; 37 cells, 4 fish; Fig. 5).

**Figure 4:**
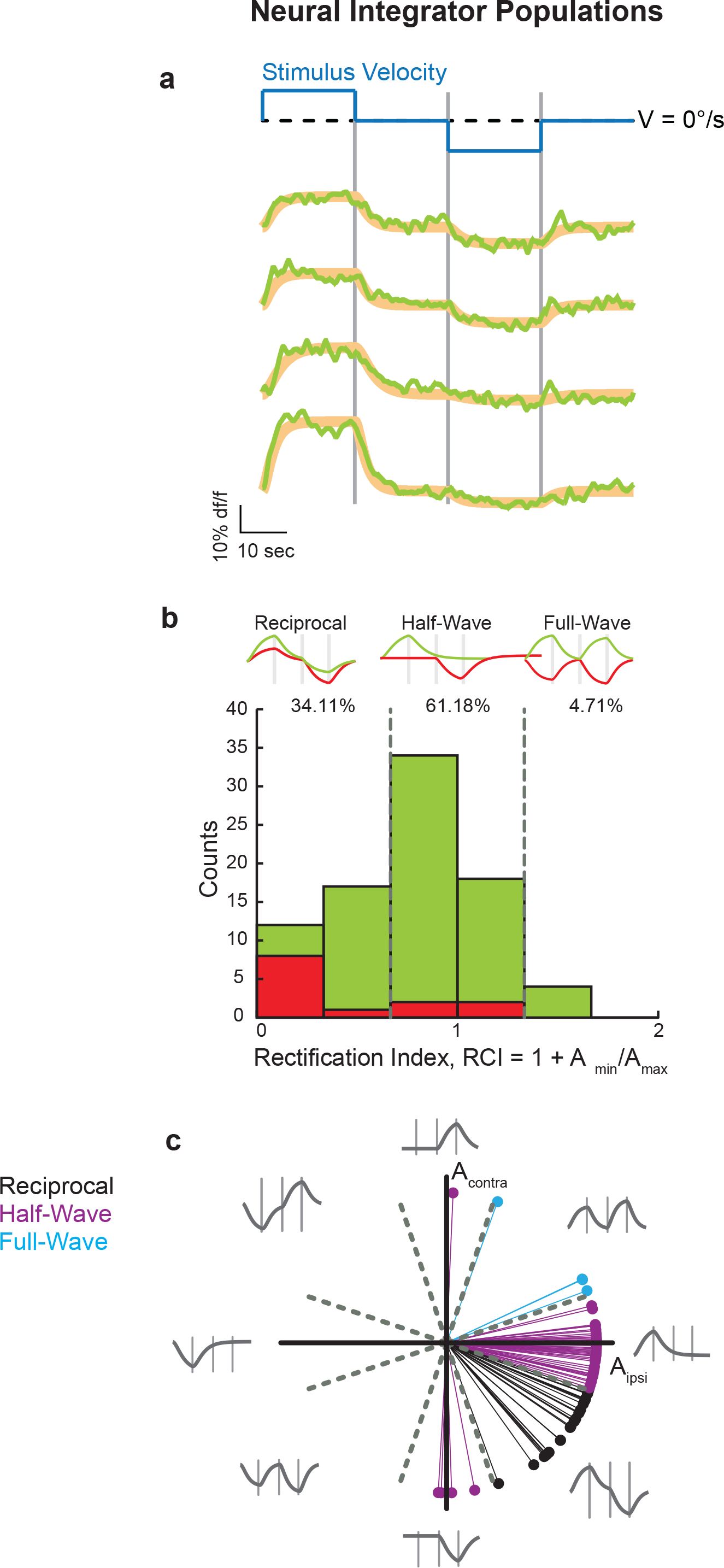
Optokinetic responses in the neural integrator populations. (**a**) Representative cycle-triggered fluorescence responses (green) of neurons in the neural integrator populations of the larval zebrafish during optokinetic stimulation (blue). Gray bars denote the time of change of stimulus velocity. Model fits are in orange.
(**b**) Histogram of the number of neurons within each response type. Arrangement as in Figure 3b. (**c**) Polar plot indicating the relative contribution of the fit coefficients Aipsi and Acontra. Grey dashed lines denote the RCI response group cutoffs in polar coordinates.

**Figure 5:**
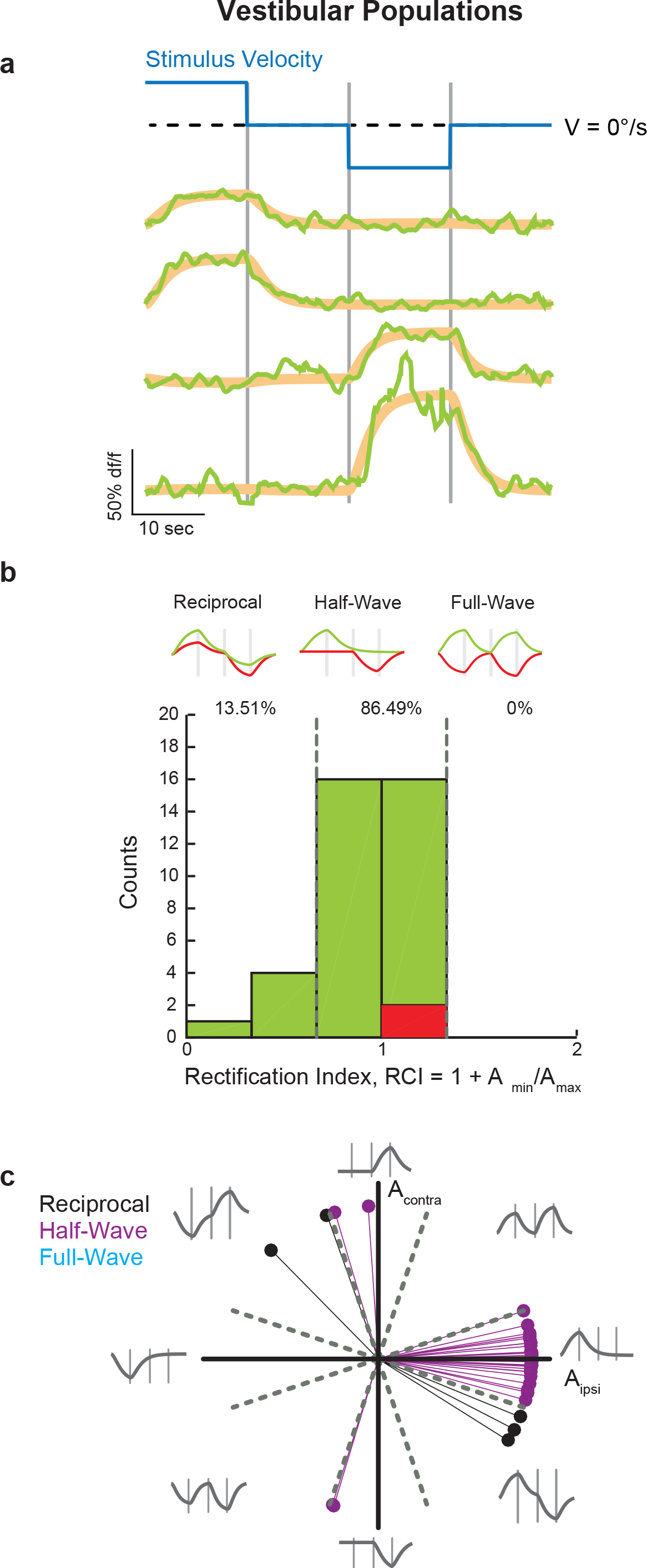
Optokinetic responses in the vestibular complex. (**a**) Representative cycle-triggered fluorescence responses (green) of four neurons in the vestibular complex of the larval zebrafish during optokinetic stimulation (blue). Gray bars denote the time of change of stimulus velocity. Model fits are in orange. (**b**) Histogram of the number of neurons within each response type Arrangement as in Figure 3b. (**c**) Polar plot indicating the relative contribution of the fit coefficients Aipsi and Acontra. Grey dashed lines denote the RCI response group cutoffs in polar coordinates.

We also looked for differences and trends associated with the dynamics of neuronal firing. The distribution of time constants *τ*_*p*_ in the granule and NI populations differed (p = 6.99×10^−5^, KS Test), with the granule cell population generally exhibiting faster relaxation dynamics (p = 4.69×10^−7^, Two-Sample T-Test): on average, the granule cell population *τ*_*p*_ was 3.94 ± 0.39 seconds (mean ± s.e.m.) while the average *p* for the NI population was 7.22 ± 0.61 seconds. The 10^th^, 50^th^, and 90^th^ percentile values of *τ*_*p*_ for the granule cell population were 0.56, 2.92, and 10.16 seconds; for the NI populations, these values were 1.32, 6.26, and 14.25 seconds, consistent with previous findings in the larval zebrafish of heterogeneity in the dynamics of VPNI neurons (Miri et al., 2011b; Daie et al., 2015) and a lack of optokinetic after-nystagmus (Beck et al., 2004). There were no notable correlations between time constant and rectification in either population (granule: p = 0.87; NI: p = 0.71; Spearman rank correlation).

The main results presented thus far can be summarized as follows. First, a sizable fraction (189/543, ~1/3) of the granule cell population was responsive during optokinetic behavior, with representations that vary in the degree of rectification and balance of excitation vs. inhibition. Second, a comparison of the patterns of activity in the granule cell and core sensorimotor populations suggests that the granule layer uses a representation that encompasses and expands upon that present in the core sensorimotor pathway.

### The spatial organization of granule cell activity

We next evaluated whether signaling in the inner granule neuron population was spatially organized. To do so, we first examined if there was any spatial dependence in the degree of functional correlation between granule cell pairs (Fig. 6). The spatial dependence of the strength of functional association between responsive granule cells was determined by calculating for every pair the correlation (or anticorrelation) between the activity time series and pairwise distance (blue). For comparison, we performed the same measurement for granule cells that were not responsive during the optokinetic behavior (black). We found that there was a weak trend for the activity time series of nearby cells to be more correlated or anti-correlated than those further away (r^2^ = 0.81, Spearman Rank Correlation, or src, −0.80; p = 2.83 × 10^−5^).

**Figure 6:**
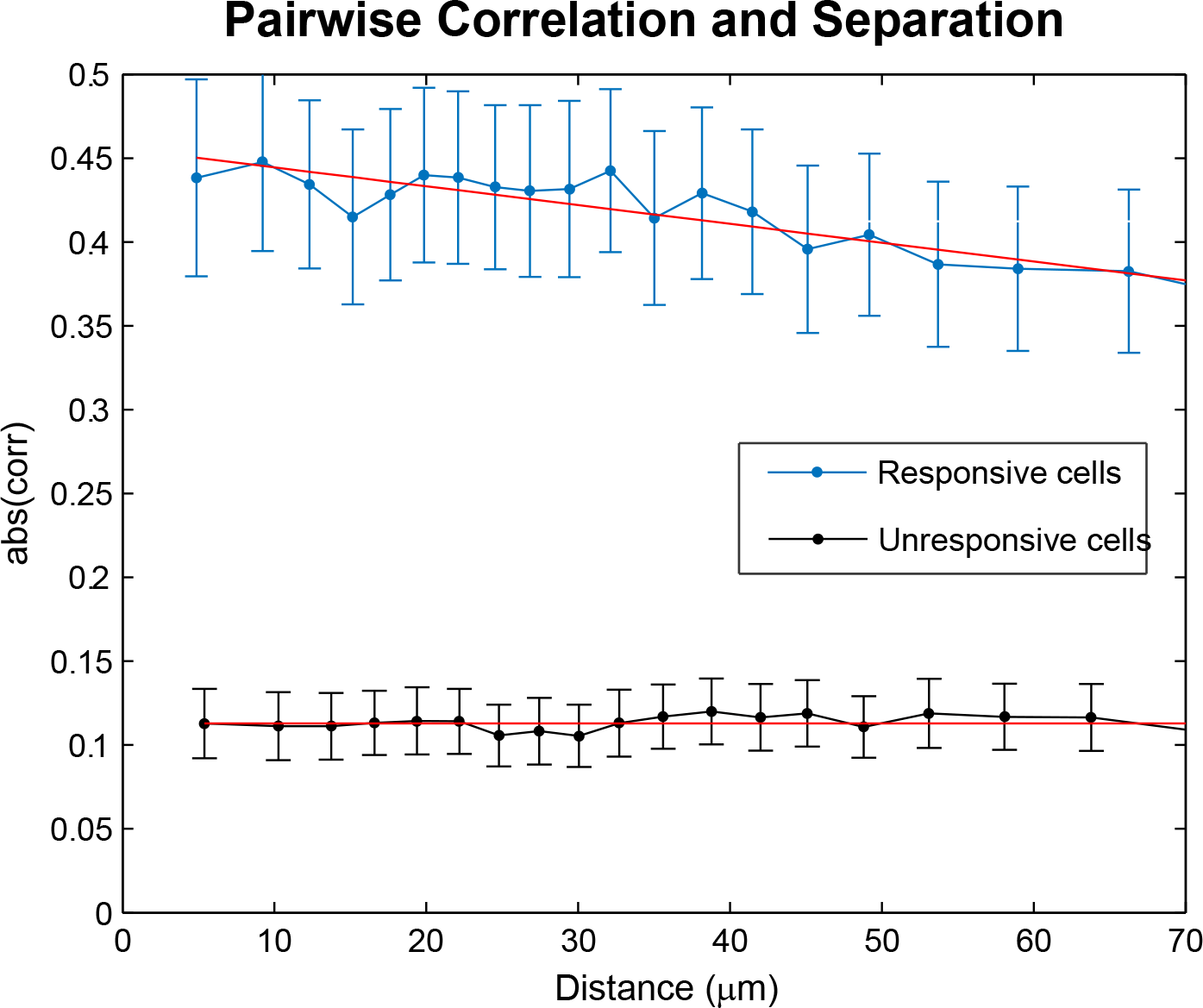
Spatial patterning in pairwise correlations. The spatial dependence of the strength of functional association between responsive granule cells was determined by calculating for every pair the correlation (or anticorrelation) between their activity time series and pairwise distance (blue). For comparison, we performed the same measurement for granule cells that were not responsive during the optokinetic behavior (black). Data were grouped into 20 bins with equal sample numbers, and then the mean +/-s.e.m. index difference was plotted against the mean separation distance for each bin. Statistics were calculated after binning.

**Figure 7:**
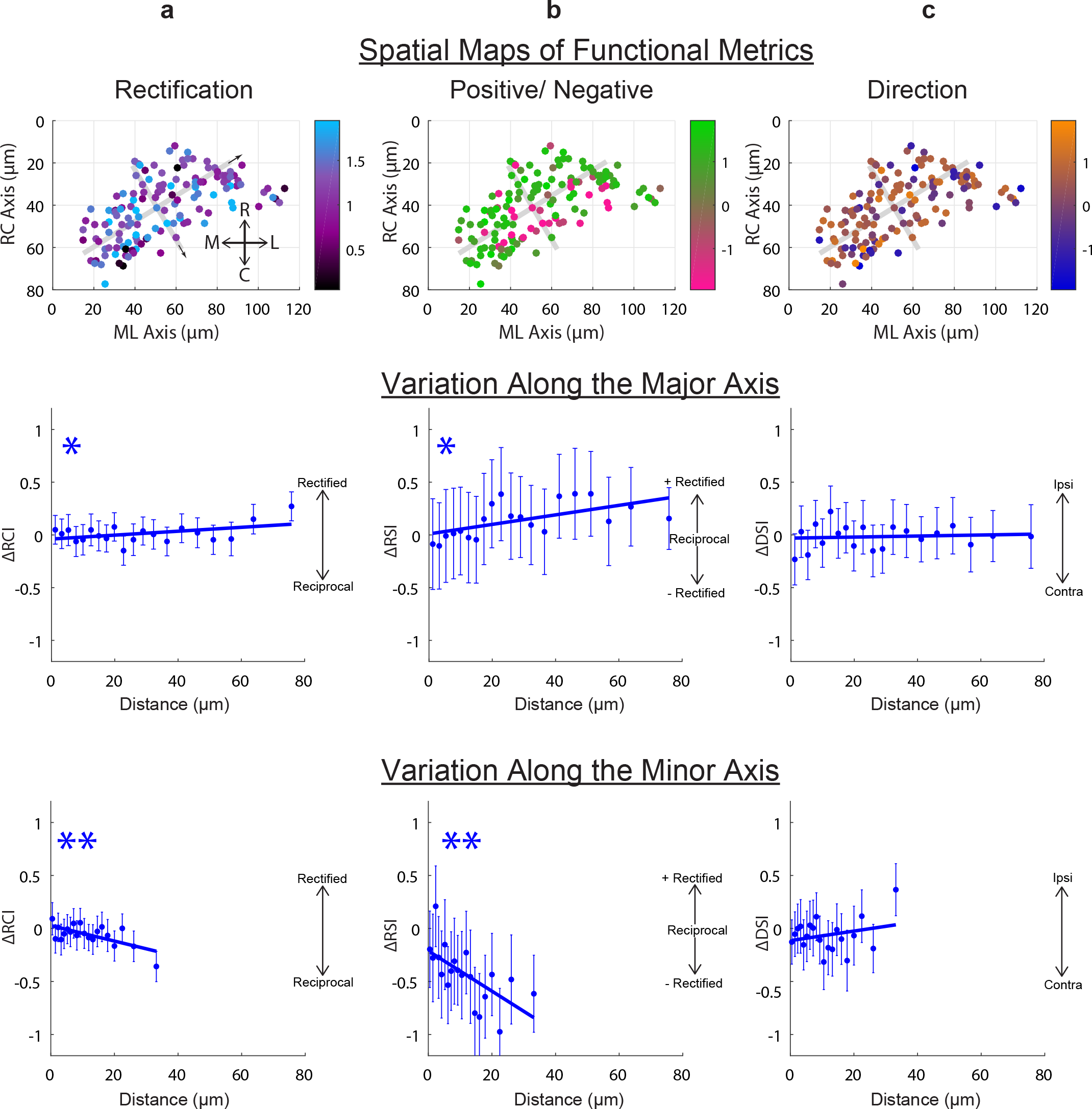
Weak spatial organization of functional properties in the granule cell population. (**a**) Spatial patterns in rectification. (Top) the cell position of each granule cell across all data sets is shown with degree of rectification color coded (data compiled onto the right lobe). Gray lines indicate the major and minor axis of the population; arrows indicate the positive direction along these axes. (Middle, bottom) Difference in rectification index (*RCI*) is plotted versus pairwise distance along the major and minor axes. Significant trends are indicate with * (p=0.05) and ** (p=0.001), and error bars are the s.e.m. of each bin. (**b**) As above, but for spatial patterns in the signed rectification index. (**c**) As in (a), but for spatial patterns in the direction sensitivity index.

Prompted by this result, we then assessed spatial organization in functional metrics (Fig. 7). Trends were assessed in two ways: first from fitting a plane to cell function and location data after registering across animals (Fig. 7, top), and second, to circumvent possible nonlinearities and registration related artifacts, by calculating pairwise differences in functional metrics and location before pooling across animals (Fig. 7, bottom). The most notable trends were found along the minor axis of the population (Table 1). We first assessed spatial organization in the degree of rectification (*RCI*, Fig. 7a). A plane of best fit exhibited a slight tilt along the minor axis of the granule population (slope = −3.2 × 10^−3^ units/μm for minor axis, 3.4 × 10 ^−4^ units/μm for major axis), but the trend did not reach significance (p>0.05). The trend along the minor axis was significant with pairwise analysis, with more rectified responses at rostro-medial locations (r^2^ = 0.42, slope = −7.64 × 10^−3^ units/μm, src = −7.52 × 10^−2^, p = 3.22 × 10^−4^). Next we looked at spatial organization in how excited or inhibited a cell was during optokinetic response using a rectification sign index (*RSI*, Fig. 7b). A plane of best fit again exhibited tilt along the minor axis (slope = −1.8 × 10^−2^ units/μm for minor axis, 2.4 × 10^−3^ units/μm for major axis) that did not achieve significance (p>0.05). This trend was significant with pairwise analysis, with more excited responses at rostro-medial locations (r^2^ = 0.40, slope = −1.86 × 10^−2^ units/μm, src = −1.18×10^−1^, p = 1.41 × 10^−8^). No significant trends were found with regards to direction sensitivity (Fig. 7c).

**Table 1:** Summary of statistics for spatial patterning in functional metrics. Summary statistics for pairwise analyses of spatial patterning in the rectification index (RCI), signed rectification index (RSI), and direction sensitivity index (DSI) along the rostro-caudal axis (RC), medio-lateral axis (ML), and major and minor axes of the granule population. Bootstrap and robust regression analysis revealed that all significant trends were robust to outliers except the trend for RSI along the ML axis.

|  |  | RCI | RSI | DSI |
| --- | --- | --- | --- | --- |
| RC axis | Slope | -8.10E-04 | -2.51E-03 | -2.34E-03 |
|  | SRC | 1.65E-02 | -1.33E-02 | -3.95E-02 |
|  | P value | 4.30E-01 | 5.25E-01 | 5.90E-02 |
| ML axis | Slope | -3.23E-03 | -3.36E-03 | 1.31E-03 |
|  | SRC | -8.41E-02 | -1.71E-01 | 3.01E-02 |
|  | P value | 5.64E-05 | 1.10E-02 | 1.51E-01 |
| Major Axis | Slope | 1.80E-03 | 4.88E-03 | 6.69E-04 |
|  | SRC | 4.17E-02 | 6.46E-02 | -1.05E-02 |
|  | P value | 4.64E-02 | 1.99E-03 | 6.15E-01 |
| Minor Axis | Slope | -7.64E-03 | -1.86E-02 | -1.38E-03 |
|  | SRC | -7.52E-02 | -1.18E-01 | 1.40E-02 |
|  | P value | 3.22E-04 | 1.41E-08 | 5.03E-01 |

Together, these data suggest that 1) there is a weak spatial organization in the functional responses of granule neurons, and 2) the strongest trends in functional properties occur along the minor axes of the population. The relationship between this direction and the patterns of inputs to and outputs from the granule layer are addressed in the Discussion.

### Implications for the cerebellar control of learning

Our experimental results suggest that the granule layer represents the information essential for learning as a set of elementary signaling motifs that emphasize those elements largely absent in the signals along the core sensorimotor pathways directly controlling behavior. We next examined the computational significance of these findings for cerebellar processing. First, we considered which patterns of activity could be obtained at the Purkinje cell layer using different combinations of the recorded granule cell subtypes (Fig. 8a). Second, we examined how the observed granule cell population coding scheme (Fig. 3c) might contribute to the efficient adaptation of Purkinje cell responses in support of motor learning (Fig. 8b).

**Figure 8:**
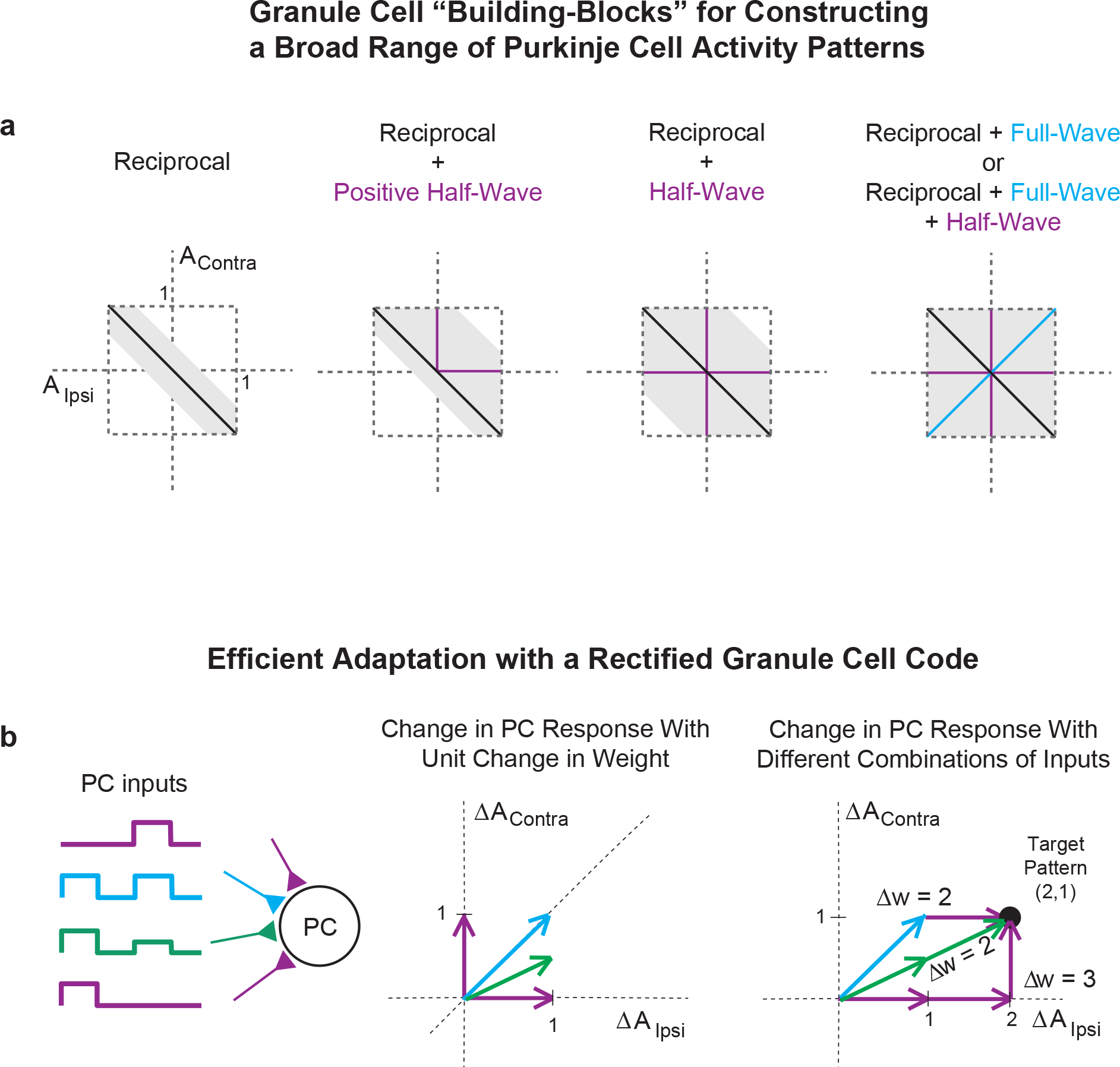
The rectified components of granule output and implications for cerebellar learning. (**a**) Possible sets of responses to ipsiversive (x-axis) and contraversive (y-axis) optokinetic stimulation for a Purkinje cell whose response reflects a weighted average of inputs from different combinations of granule cell subtypes. Columns: Possible Purkinje cells responses if only reciprocal granule neuron responses are available (1^st^ column), or if, additionally, positive half-wave rectified (2^nd^ column, assuming granule cells drives net excitation), all half-wave rectified (or, equivalently, positive half-wave rectified that are able to drive net inhibition) (3^rd^ column), or full-wave rectified granule cells (4^th^ column) are available. Individual granule cell responses were allowed to occupy angles of +/− 15 degrees around the idealized directions (black, purple, or cyan), and were constrained to have a maximal x or y coordinate of 1. (**b**) Efficiency of learning with different types of granule cells. Left, four example granule neuron inputs onto a Purkinje cell (PC). Middle, change in PC response to ipsilateral (x-axis) or contralateral (y-axis) velocity stimuli following a 1-unit change in weight from each of the shown granule cell types. Each granule cell was assumed to have a response of 1 unit amplitude to ipsilateral or contralateral stimuli, leading to the shown changes in PC firing (arrows) if the corresponding input’s weight was changed. Note that granule cells which are active in response to both inputs produce larger vector-amplitude changes in PC response. Right, three possible sets of weight changes leading to a change (2,1) in PC response. Use of only half-wave rectified cells (purple) requires 3 units of weight change (3 purple arrows). Changes in either granule cells with the desired ratio of ipsilateral-to-contralateral sensitivity (2 green arrows), or use of full-wave rectified cells (cyan arrow), requires only 2 units of weight change.

We explored these questions using a phenomenological model in which both granule and Purkinje neuron responses were characterized by a vector **v** = (*A*_*ipsi*_, *A*_*contra*_) giving their response amplitudes to ipsilateral and contralateral stimulus velocity. Granule neuron responses to ipsilateral or contralateral movements were allowed to vary in amplitude from −1 to 1 unit. Purkinje cell activity was generated by a weighted sum of the individual granule neuron responses 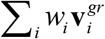, where *w*_*i*_ is the weight (positive) from the *i*^th^ granule cell and the total weight onto a Purkinje cell was constrained such that ∑_*i*_ *w*_*i*_ ≤ 1.

Figure 8a shows the possible patterns of Purkinje cell activity that could be achieved by summing different subpopulations of the recorded granule cell responses. Reciprocal granule cells exhibit strong negative correlations between their response amplitudes to ipsiversive and contraversive stimuli. As a result, for a fixed maximal amplitude of individual granule-to-Purkinje neuron weights, only a narrow band of Purkinje cell responses is possible when using these inputs alone (Fig. 8a, reciprocal). If excitatory input from positive half-wave rectified neurons also is available, then a much larger portion of the possible space of responses can be explored (Fig. 8a, reciprocal + positive half-wave). This space is expanded further if both positive and negative half-wave rectified granule neurons also are used (Fig. 8a, reciprocal + half-wave), or equivalently if positive half-wave rectified cells are able to drive inhibition of Purkinje cells through disynaptic inputs through stellate or basket cells (Barmack and Yakhnitsa, 2008; Dean et al., 2010). Finally, the complete response space can be explored if full-wave rectified neurons are also available (Fig. 8a, far right). In summary, we see that the presence of rectified signaling components at the granule cell layer enables the cerebellum to generate a wide array of Purkinje cell responses to facilitate adaptive behavior.

We next asked how the recorded distribution of granule cell responses, characterized by a broad heterogeneity and over-representation of full-wave rectified activity patterns (Fig. 3c), might contribute to the efficient adaptation of Purkinje cell responses (Fig. 8b). To address this question, we computed the total synaptic weight change onto a Purkinje cell required to achieve a given change in Purkinje cell response for different combinations of granule cell inputs. In Fig. 8b, we consider for illustration the case in which the desired change in Purkinje cell response is given by the target pattern (*ΔA*_*ipsi*_, *ΔA*_*contra*_) = (2,1). Granule neurons were normalized to have individual maximal responses to ipsiversive or contraversive movement of unit amplitude (Fig. 8b, middle). Half-wave rectified neurons alone could be used to achieve the target pattern, but this strategy required 3 units of weight change (Fig. 8b, right, purple path, individual arrows indicate effect of 1 unit weight change at the synapse from the shown neuronal type). By contrast, using granule neurons with response patterns matching the desired pattern change required only two units of weight change (green path). Interestingly, this same minimal weight change can be accomplished by using 1 unit of weight change along a full-wave rectified cell (cyan) to accomplish the portion of the target pattern involving both *A*_*ipsi*_ and *A*_*contra*_, plus one unit of weight change along a half-rectified neuron (top, purple arrow) to reach the desired target pattern. Together, these results suggest a role both for a diversity of granule neuron responses that individually match any given target direction, and a particular role for full-wave rectified granule neurons in rapidly altering activity along multiple directions simultaneously.

## DISCUSSION

Our data provide one of the first looks at coding properties throughout a cerebellar granule cell population during behavior. In the following, we first address the differences between the activity of granule cells and those of neurons in the core sensorimotor populations controlling optokinetic behavior. Second, we discuss our finding of a weak spatial organization of functional properties. Third we consider the implications of these results for models of cerebellar function and learning.

Granule neuron activity patterns differed from those in core sensorimotor pathway in several ways. As presented in the introduction, the core sensorimotor regions involved in the optokinetic response are the pretectum, vestibular nuclei, and the velocity-storage (VSNI) and velocity-to-position neural integrators (VPNI) (Mustari et al., 1994; Buttner-Ennever and Horn, 1997; Beck et al., 2006; Straka et al., 2006; Masseck and Hoffmann, 2009b; Portugues et al., 2014). These regions are all a source of optokinetic signals to the cerebellum (Finger, 1978; Langer et al., 1985; Wullimann and Northcutt, 1988; Mustari et al., 1994; Pastor et al., 1994; Straka et al., 2006; Ma et al., 2009; Masseck and Hoffmann, 2009b; Kolkman et al., 2011; Lee et al., 2015). Recordings in these regions in fish and mammals reveal reciprocal and positive half-wave signals during back-and-forth whole-field optokinetic stimulation (Soodak and Simpson, 1988; Mustari and Fuchs, 1990; Pastor et al., 1994; Stahl and Simpson, 1995; Beck et al., 2006; Masseck and Hoffmann, 2009a; Kubo et al., 2014; Portugues et al., 2014; Lee et al., 2015). Furthermore, recordings during oculomotor tasks of mossy fibers in the primate also reveal reciprocal or positively half-wave rectified signals (Lisberger and Fuchs, 1978; Miles et al., 1980; Noda and Warabi, 1987; Prsa et al., 2009). Our recordings in the two NI populations and the vestibular complex confirmed these previous observations that the core sensorimotor populations carry reciprocal and positive half-wave rectified responses during optokinetic tracking. At the inner granule cell population, however, in addition to these expected waveforms, we found negative half-wave rectified responses, and both positive and negative full-wave rectified responses.

One possibility for the difference could be mossy fiber input from sources not typically associated with optokinetic behavior. A potential source could be the optic tectum. Previous imaging studies have reported whole-field, movement sensitive activity in the neuropil (Portugues et al., 2014) and superficial cell body regions (Niell and Smith, 2005) of the tectum that we would categorize as positive, full-wave rectified responses. However, anatomical studies have not identified tecto-cerebellar projections (Finger, 1978; Luiten, 1981; Wullimann and Northcutt, 1988), and tectal ablations do not impact optokinetic behavior (Springer et al., 1977; Roeser and Baier, 2003). Another potential source of rectified inputs could be proprioceptive signals from the extraocular muscles. Single unit recordings from Purkinje cells in the cerebellum during passive ocular deflections or extraocular muscle stretch reveal a spectrum of responses, including full-wave rectified and inhibitory (Schwarz and Tomlinson, 1977; Ashton et al., 1989). However, recordings of the primary afferents carrying proprioceptive information show reciprocal responses (Fahy and Donaldson, 1998), although the sampling to date has been rather small (Donaldson, 2000). Both of these scenarios warrant further consideration.

A straightforward scenario for how the varied responses we observed in the granule layer could arise involves a recoding of mossy fiber inputs from the core sensorimotor pathways. Recent work in a cerebellar-like structure of the electric fish (Kennedy et al., 2014) suggests that granule cell responses might be simply explained by random combinations of mossy fiber inputs. While some features of granule cell responses would naturally emerge from the combinatorics of randomly combining half-wave rectified and reciprocal cells, it is less clear whether such random additive combinations could explain all features of our recorded granule cell populations. Direct mossy-fiber input would most simply explain granule neurons exhibiting reciprocal waveforms or positive half-wave rectification. Positive full-wave rectification, on the other hand, could arise by combining inputs from mossy fibers with opposing patterns of firing; for example, the combination of input from ipsilaterally and contralaterally located VSNI neurons exhibiting positive half-wave rectification (Straka et al., 2006) would result in exclusively increased firing during stimulation. Negatively rectified patterns would likely require intra-cerebellar processing since the vast majority of mossy fibers are excitatory (Chan-Palay, 1982; Hamori and Takacs, 1989) and, to our knowledge, negatively rectified responses during optokinetic behavior in the core sensorimotor populations have not been reported. This intra-cerebellar processing could involve metabotropic regulation of potassium currents at unipolar brush cells (Borges-Merjane and Trussell, 2015), recently identified Purkinje cell collaterals onto granule cells (Guo et al., 2016) or pathways that include Golgi cells (Medina et al., 2000; Chadderton et al., 2004; Bengtsson et al., 2012).

We also found weak spatial organization in some of the functional properties of the granule population. This suggests a non-random pattern of functional inputs to the Purkinje layer, in agreement with recent recordings in anesthetized mice of parallel fiber activity during sensory stimulation (Wilms and Hausser, 2015). The potential utility of the organization is open to debate, but considerations should include redundancy for increases the signal-to-noise ratio of the inputs to the Purkinje cell (Spanne and Jorntell, 2013). Interestingly, the strongest functional gradients were aligned with the minor axis of the inner granule cell population, an orientation that is normal to the mossy fiber input projections. This is consistent with the idea that the lack of randomness observed in granule cell activity patterns may derive from local organization in the divergence of mossy fiber inputs onto this population.

The differences in the activity patterns between the core sensorimotor populations and the granule cell population inform debates in the field about how cerebellar computations may be involved in learning. Theories of cerebellar processing have proposed that the granule layer of the cerebellum recodes its inputs into an expanded representation facilitating learning (Marr, 1969; Albus, 1971; Sejnowski, 1977; Fujita, 1982; Dean and Porrill, 2008), and support for expansion recoding has been found in electric fish (Sawtell, 2010; Kennedy et al., 2014) and rodents (Huang et al., 2013; Chabrol et al., 2015; Ishikawa et al., 2015). Expansion recoding theories have differed in the proposed degree of expansion occurring in the granule cell layer. In the earliest theories, each granule cell coded for distinct combinations of sensorimotor parameters at any given moment, or even different values of a single parameter, leading to an extremely sparse representation of cerebellar inputs (Marr, 1969; Albus, 1971). Later work instead postulated that groups of granule neurons encode elementary features of the mossy-fiber inputs, allowing sensorimotor signals to be efficiently represented through a combination of basis functions or ‘filters’ (Sejnowski, 1977; Fujita, 1982; Dean and Porrill, 2008). Our recordings over a broad swath (>75%) of a granule population during simple oculomotor behavior showed that responses did not appear sparse either in space or time, which complements recent findings in zebrafish (Knogler et al., 2017) and mice(Ozden et al., 2012; Powell et al., 2015) that suggest granule cells also use a non-sparse code during more complex locomotor behaviors. The lack of sparsity we observed seems most consistent with the idea that, at least during oculomotor learning behaviors in the larval zebrafish like VPNI plasticity (Major et al., 2004), adaptations are supported by a relatively limited expansion of sensorimotor signals into a set of elementary motifs (Sejnowski, 1977; Fujita, 1982; Dean and Porrill, 2008; Spanne and Jorntell, 2013). Future work will be needed to see if this representational scheme changes dramatically over the course of development as the number of cells in the zebrafish cerebellum expands or is complemented by sparsity on finer time scales not easily accessible to calcium imaging.

A separate debate in the cerebellar field has focused on whether the site (or sites) of plasticity underlying motor adaptations reside in the cerebellar cortex, or, alternatively, if the cerebellar cortex serves as a teacher instructing plasticity in core sensorimotor populations (Raymond et al., 1996; Ito, 2006). Our data provide a possible alternative viewpoint that contains elements of each hypothesis (Boyden et al., 2006; Katoh et al., 2008). While full-wave rectified responses were highly represented in the granule neuron population recorded, reciprocal responses prevalent in the core populations mediating optokinetic behavior were least well represented. Hence, while modifications such as changes in the gain of the vestibulo-ocular reflex could be achieved simply through adaptation at non-cerebellar locations in the strength of reciprocal signals, the signals in the granule population appear best-suited to mediate non-reciprocal changes in behavior, for example in compensation for an injury at the oculomotor plant that necessitates an asymmetric adaptation in the neural drive to an agonist/antagonist muscle pair (Hirata et al., 2012). More generally, whereas simple learning behaviors like gain adaptation may require cerebellar activity only to instruct downstream sites of plasticity, learning that involves movements along directions poorly encoded in the core sensorimotor populations may depend critically on plasticity within the cerebellar cortex.

## Acknowledgements

The authors thank Robert Baker, Tom Chartrand, Kayvon Daie, Jeremy Dittman, Andrea Giovannucci, Masahiko Hibi, Jennifer Raymond, Takashi Shimizu, Jerry Simpson, David Tank, Jonathan Victor, and Sam Wang for helpful comments on this work, and Kayvon Daie for assistance with data analysis. Funding for this work was provided by NIH Training grant EY007138-16 (SS); the Burroughs Wellcome Career Award at the Scientific Interface (EA); a UC Davis Ophthalmology Research to Prevent Blindness grant (MG); a Simons Collaboration on the Global Brain grant, and NIH grant R01 EY021581 (EA and MG).

## REFERENCES

Aksay, E, Olasagasti, I, Mensh, BD, Baker, R, Goldman, MS, Tank DW (2007) Functional dissection of circuitry in a neural integrator. Nat Neurosci 10:494–504.

Albus J (1971) A theory of cerebellar function. Math Biosci 10:25–61.

Ashton, JA, Milleret, C, Donaldson IM (1989) Effects of afferent signals from the extraocular muscles upon units in the cerebellum, vestibular nuclear complex and oculomotor nucleus of the trout. Neuroscience 31:529–541.

Bae, YK, Kani, S, Shimizu, T, Tanabe, K, Nojima, H, Kimura, Y, Higashijima, S, Hibi M (2009) Anatomy of zebrafish cerebellum and screen for mutations affecting its development. Dev Biol 330:406–426.

Bahn, S, Harvey, RJ, Darlison, MG, Wisden W (1996) Conservation of gamma-aminobutyric acid type A receptor alpha 6 subunit gene expression in cerebellar granule cells. Journal of neurochemistry 66:1810–1818.

Barmack, NH, Yakhnitsa V (2008) Functions of interneurons in mouse cerebellum. J Neurosci 28:1140–1152.

Beck, JC, Gilland, E, Tank, DW, Baker R (2004) Quantifying the ontogeny of optokinetic and vestibuloocular behaviors in zebrafish, medaka, and goldfish. J Neurophysiol 92:3546–3561.

Beck, JC, Rothnie, P, Straka, H, Wearne, SL, Baker R (2006) Precerebellar hindbrain neurons encoding eye velocity during vestibular and optokinetic behavior in the goldfish. J Neurophysiol 96:1370–1382.

Bengtsson, F, Geborek, P, Jorntell H (2012) Cross-correlations between pairs of neurons in cerebellar cortex in vivo. Neural networks: the official journal of the International Neural Network Society.

Borges-Merjane C, Trussell LO (2015) ON and OFF unipolar brush cells transform multisensory inputs to the auditory system. Neuron 85:1029–1042.

Boyden, ES, Katoh, A, Pyle, JL, Chatila, TA, Tsien, RW, Raymond JL (2006) Selective engagement of plasticity mechanisms for motor memory storage. Neuron 51:823–834.

Brondi, M, Sato, SS, Rossi, LF, Ferrara, S, Ratto GM (2012) Finding a Needle in a Haystack: Identification of EGFP Tagged Neurons during Calcium Imaging by Means of Two-Photon Spectral Separation. Frontiers in molecular neuroscience 5:96.

Buttner-Ennever JA, Horn AK (1997) Anatomical substrates of oculomotor control. Current opinion in neurobiology 7:872–879.

Chabrol, FP, Arenz, A, Wiechert, MT, Margrie, TW, DiGregorio DA (2015) Synaptic diversity enables temporal coding of coincident multisensory inputs in single neurons. Nat Neurosci 18:718–727.

Chadderton, P, Margrie, TW, Hausser M (2004) Integration of quanta in cerebellar granule cells during sensory processing. Nature 428:856–860.

Chan-Palay V (1982) Gamma-aminobutyric acid pathways in the cerebellum studied by retrograde and anterograde transport of glutamic acid decarboxylase (GAD) antibody after in vivo injections. Progress in brain research 55:51–76.

D'Angelo, E, Rossi, P, Taglietti V (1994) Voltage-dependent kinetics of N-methyl-D-aspartate synaptic currents in rat cerebellar granule cells. The European journal of neuroscience 6:640–645.

D'Angelo, E, De Filippi G, Rossi, P, Taglietti V (1998) Ionic mechanism of electroresponsiveness in cerebellar granule cells implicates the action of a persistent sodium current. J Neurophysiol 80:493–503.

Daie, K, Goldman, MS, Aksay ER (2015) Spatial patterns of persistent neural activity vary with the behavioral context of short-term memory. Neuron 85:847–860.

Dean, P, Porrill J (2008) Adaptive-filter models of the cerebellum: computational analysis. Cerebellum 7:567–571.

Dean, P, Porrill, J, Ekerot, CF, Jorntell H (2010) The cerebellar microcircuit as an adaptive filter: experimental and computational evidence. Nat Rev Neurosci 11:30–43.

Dino, MR, Mugnaini E (2008) Distribution and phenotypes of unipolar brush cells in relation to the granule cell system of the rat cochlear nucleus. Neuroscience 154:29–50.

Donaldson IM (2000) The functions of the proprioceptors of the eye muscles. Philosophical transactions of the Royal Society of London Series, B, Biological sciences 355:1685–1754.

Dunn, TW, Mu, Y, Narayan, S, Randlett, O, Naumann, EA, Yang, CT, Schier, AF, Freeman, J, Engert, F, Ahrens MB (2016) Brain-wide mapping of neural activity controlling zebrafish exploratory locomotion. eLife 5:e12741.

Fahy, FL, Donaldson IM (1998) Signals of eye position and velocity in the first-order afferents from pigeon extraocular muscles. Vision research 38:1795–1804.

Finger TE (1978) Cerebellar afferents in teleost catfish (Ictaluridae). The Journal of comparative neurology 181:173–181.

Fujita M (1982) Adaptive filter model of the cerebellum. Biol Cybern 45:195–206.

Gall, D, Prestori, F, Sola, E, D'Errico, A, Roussel, C, Forti, L, Rossi, P, D'Angelo E (2005) Intracellular calcium regulation by burst discharge determines bidirectional long-term synaptic plasticity at the cerebellum input stage. J Neurosci 25:4813–4822.

Gandolfi, D, Pozzi, P, Tognolina, M, Chirico, G, Mapelli, J, D'Angelo E (2014) The spatiotemporal organization of cerebellar network activity resolved by two-photon imaging of multiple single neurons. Frontiers in cellular neuroscience 8:92.

Guo, C, Witter, L, Rudolph, S, Elliott, HL, Ennis, KA, Regehr WG (2016) Purkinje Cells Directly Inhibit Granule Cells in Specialized Regions of the Cerebellar Cortex. Neuron 91:1330–1341.

Hamling, KR, Tobias, ZJ, Weissman TA (2015) Mapping the development of cerebellar Purkinje cells in zebrafish. Developmental neurobiology 75:1174–1188.

Hamori, J, Takacs J (1989) Two types of GABA-containing axon terminals in cerebellar glomeruli of cat: an immunogold-EM study. Experimental brain research Experimentelle Hirnforschung Experimentation cerebrale 74:471–479.

Hirata, Y, Katagiri, K, Tanaka Y (2012) Direct causality between single-Purkinje cell activities and motor learning revealed by a cerebellum-machine interface utilizing VOR adaptation paradigm. Cerebellum 11:455–456.

Huang, CC, Sugino, K, Shima, Y, Guo, C, Bai, S, Mensh, BD, Nelson, SB, Hantman AW (2013) Convergence of pontine and proprioceptive streams onto multimodal cerebellar granule cells. eLife 2:e00400.

Ishikawa, T, Shimuta, M, Hausser M (2015) Multimodal sensory integration in single cerebellar granule cells in vivo. eLife 4.

Ito M (2006) Cerebellar circuitry as a neuronal machine. Progress in neurobiology 78:272–303.

Jorntell, H, Hansel C (2006) Synaptic memories upside down: bidirectional plasticity at cerebellar parallel fiber-Purkinje cell synapses. Neuron 52:227–238.

Kaslin, J, Kroehne, V, Benato, F, Argenton, F, Brand M (2013) Development and specification of cerebellar stem and progenitor cells in zebrafish: from embryo to adult. Neural development 8:9.

Kato K (1990) Novel GABAA receptor alpha subunit is expressed only in cerebellar granule cells. Journal of molecular biology 214:619–624.

Katoh, A, Chapman, PJ, Raymond JL (2008) Disruption of learned timing in P/Q calcium channel mutants. PLoS One 3:e3635.

Kennedy, A, Wayne, G, Kaifosh, P, Alvina, K, Abbott, LF, Sawtell NB (2014) A temporal basis for predicting the sensory consequences of motor commands in an electric fish. Nat Neurosci 17:416–422.

Kinkhabwala, A, Riley, M, Koyama, M, Monen, J, Satou, C, Kimura, Y, Higashijima, S, Fetcho J (2011) A structural and functional ground plan for neurons in the hindbrain of zebrafish. Proc Natl Acad Sci U S A 108:1164–1169.

Knogler, LD, Markov, DA, Dragomir, EI, Stih, V, Portugues R (2017) Sensorimotor Representations in Cerebellar Granule Cells in Larval Zebrafish Are, Dense, Spatially, Organized, and Non-temporally Patterned. Current biology: CB 27:1288–1302.

Kolkman, KE, McElvain, LE, du Lac S (2011) Diverse precerebellar neurons share similar intrinsic excitability. J Neurosci 31:16665–16674.

Kubo, F, Hablitzel, B, Dal Maschio M, Driever, W, Baier, H, Arrenberg AB (2014) Functional architecture of an optic flow-responsive area that drives horizontal eye movements in zebrafish. Neuron 81:1344–1359.

Langer, T, Fuchs, AF, Scudder, CA, Chubb MC (1985) Afferents to the flocculus of the cerebellum in the rhesus macaque as revealed by retrograde transport of horseradish peroxidase. The Journal of comparative neurology 235:1–25.

Lee, MM, Arrenberg, AB, Aksay ER (2015) A structural and genotypic scaffold underlying temporal integration. J Neurosci 35:7903–7920.

Lisberger, SG, Fuchs AF (1978) Role of primate flocculus during rapid behavioral modification of vestibuloocular reflex. II. Mossy fiber firing patterns during horizontal head rotation and eye movement. J Neurophysiol 41:764–777.

Lopez-Barneo J, Darlot, C, Berthoz, A, Baker R (1982) Neuronal activity in prepositus nucleus correlated with eye movement in the alert cat. J Neurophysiol 47:329–352.

Luiten PG (1981) Afferent and efferent connections of the optic tectum in the carp (Cyprinus carpio L.). Brain Res 220:51–65.

Ma, LH, Punnamoottil, B, Rinkwitz, S, Baker R (2009) Mosaic hoxb4a neuronal pleiotropism in zebrafish caudal hindbrain. PLoS One 4:e5944.

Major, G, Baker, R, Aksay, E, Mensh, B, Seung, HS, Tank DW (2004) Plasticity and tuning by visual feedback of the stability of a neural integrator. Proc Natl Acad Sci U S A 101:7739–7744.

Marr D (1969) A theory of cerebellar cortex. J Physiol 202:437–470.

Masseck, OA, Hoffmann KP (2009a) Question of reference frames: visual direction-selective neurons in the accessory optic system of goldfish. J Neurophysiol 102:2781–2789.

Masseck, OA, Hoffmann KP (2009b) Comparative neurobiology of the optokinetic reflex. Ann N Y Acad Sci 1164:430–439.

Matsui, H, Namikawa, K, Babaryka, A, Koster RW (2014) Functional regionalization of the teleost cerebellum analyzed in vivo. Proc Natl Acad Sci U S A 111:11846–11851.

McFarland, JL, Fuchs AF (1992) Discharge patterns in nucleus prepositus hypoglossi and adjacent medial vestibular nucleus during horizontal eye movement in behaving macaques. J Neurophysiol 68:319–332.

Medina, JF, Garcia, KS, Nores, WL, Taylor, NM, Mauk MD (2000) Timing mechanisms in the cerebellum: testing predictions of a large-scale computer simulation. J Neurosci 20:5516–5525.

Miles, FA, Fuller, JH, Braitman, DJ, Dow BM (1980) Long-term adaptive changes in primate vestibuloocular reflex. III. Electrophysiological observations in flocculus of normal monkeys. J Neurophysiol 43:1437–1476.

Miri, A, Daie, K, Burdine, RD, Aksay, E, Tank DW (2011a) Regression-based identification of behavior-encoding neurons during large-scale optical imaging of neural activity at cellular resolution. J Neurophysiol 105:964–980.

Miri, A, Daie, K, Arrenberg, AB, Baier, H, Aksay, E, Tank DW (2011b) Spatial gradients and multidimensional dynamics in a neural integrator circuit. Nature Neuroscience 14:1150–1159.

Mustari, MJ, Fuchs AF (1990) Discharge patterns of neurons in the pretectal nucleus of the optic tract (NOT) in the behaving primate. J Neurophysiol 64:77–90.

Mustari, MJ, Fuchs, AF, Kaneko, CR, Robinson FR (1994) Anatomical connections of the primate pretectal nucleus of the optic tract. The Journal of comparative neurology 349:111–128.

Niell, CM, Smith SJ (2005) Functional imaging reveals rapid development of visual response properties in the zebrafish tectum. Neuron 45:941–951.

Nieuwenhuys R (1967) Comparative anatomy of the cerebellum. Progress in brain research 25:1–93.

Noda, H, Warabi T (1987) Responses of Purkinje cells and mossy fibres in the flocculus of the monkey during sinusoidal movements of a visual pattern. J Physiol (Lond) 387:611–628.

Ozden, I, Dombeck, DA, Hoogland, TM, Tank, DW, Wang SSH (2012) Widespread State-Dependent Shifts in Cerebellar Activity in Locomoting Mice. PLoS One 7.

Pastor, AM, De la Cruz, RR, Baker R (1994) Eye position and eye velocity integrators reside in separate brainstem nuclei. Proc Natl Acad Sci U S A 91:807–811.

Pologruto, TA, Sabatini, BL, Svoboda K (2003) ScanImage: Flexible software for operating laser scanning microscopes. BioMedical Engineering OnLine 2.

Portugues, R, Feierstein, CE, Engert, F, Orger MB (2014) Whole-brain activity maps reveal stereotyped, distributed networks for visuomotor behavior. Neuron 81:1328–1343.

Powell, K, Mathy, A, Duguid, I, Hausser M (2015) Synaptic representation of locomotion in single cerebellar granule cells. eLife 4.

Prsa, M, Dash, S, Catz, N, Dicke, PW, Thier P (2009) Characteristics of responses of Golgi cells and mossy fibers to eye saccades and saccadic adaptation recorded from the posterior vermis of the cerebellum. J Neurosci 29:250–262.

Raymond, JL, Lisberger, SG, Mauk MD (1996) The cerebellum: a neuronal learning machine& Science 272:1126–1131.

Roeser, T, Baier H (2003) Visuomotor behaviors in larval zebrafish after GFP-guided laser ablation of the optic tectum. J Neurosci 23:3726–3734.

Sawtell NB (2010) Multimodal integration in granule cells as a basis for associative plasticity and sensory prediction in a cerebellum-like circuit. Neuron 66:573–584.

Schwartz, EJ, Rothman, JS, Dugue, GP, Diana, M, Rousseau, C, Silver, RA, Dieudonne S (2012) NMDA receptors with incomplete Mg(2)(+) block enable low-frequency transmission through the cerebellar cortex. J Neurosci 32:6878–6893.

Schwarz, DW, Tomlinson RD (1977) Neuronal responses to eye muscle stretch in cerebellar lobule VI of the cat. Experimental brain research Experimentelle Hirnforschung Experimentation cerebrale 27:101–111.

Sejnowski TJ (1977) Storing covariance with nonlinearly interacting neurons. Journal of mathematical biology 4:303–321.

Soodak, RE, Simpson JI (1988) The accessory optic system of rabbit. I. Basic visual response properties. J Neurophysiol 60:2037–2054.

Spanne, A, Jorntell H (2013) Processing of Multi-dimensional Sensorimotor Information in the Spinal and Cerebellar Neuronal Circuitry: A New Hypothesis. PLoS computational biology 9:e1002979.

Springer, AD, Easter, SS Jr., Agranoff BW (1977) The role of the optic tectum in various visually mediated behaviors of goldfish. Brain Res 128:393–404.

Stahl, JS, Simpson JI (1995) Dynamics of rabbit vestibular nucleus neurons and the influence of the flocculus. J Neurophysiol 73:1396–1413.

Straka, H, Baker, R, Gilland E (2001) Rhombomeric organization of vestibular pathways in larval frogs. The Journal of comparative neurology 437:42–55.

Straka, H, Beck, JC, Pastor, AM, Baker R (2006) Morphology and physiology of the cerebellar vestibulolateral lobe pathways linked to oculomotor function in the goldfish. J Neurophysiol 96:1963–1980.

Strick, PL, Dum, RP, Fiez JA (2009) Cerebellum and nonmotor function. Annu Rev Neurosci 32:413–434.

Volkmann, K, Rieger, S, Babaryka, A, Koster RW (2008) The zebrafish cerebellar rhombic lip is spatially patterned in producing granule cell populations of different functional compartments. Dev Biol 313:167–180.

Wilms, CD, Hausser M (2015) Reading out a spatiotemporal population code by imaging neighbouring parallel fibre axons in vivo. Nature communications 6:6464.

Wullimann, MF, Northcutt RG (1988) Connections of the corpus cerebelli in the green sunfish and the common goldfish: a comparison of perciform and cypriniform teleosts. Brain, behavior and evolution 32:293–316.

Yaksi, E, Friedrich RW (2006) Reconstruction of firing rate changes across neuronal populations by temporally deconvolved Ca2+ imaging. Nat Methods 3:377–383.

